# High-throughput sequencing of macaque basolateral amygdala projections reveals dissociable connectional motifs with frontal cortex

**DOI:** 10.1101/2023.01.18.524407

**Authors:** Zachary R Zeisler, Liza London, William G Janssen, J Megan Fredericks, Catherine Elorette, Atsushi Fujimoto, Huiqing Zhan, Brian E Russ, Roger L Clem, Patrick R Hof, Frederic M Stoll, Peter H Rudebeck

## Abstract

The basolateral amygdala (BLA) projects widely across the macaque frontal cortex^1–4^, and amygdalo-frontal projections are critical for optimal emotional responding^5^ and decision-making^6^. Yet, little is known about the single-neuron architecture of these projections: namely, whether single BLA neurons project to multiple parts of the frontal cortex. Here, we use MAPseq^7^ to determine the projection patterns of over 3000 macaque BLA neurons. We found that one-third of BLA neurons have two or more distinct targets in parts of frontal cortex and of subcortical structures. Further, we reveal non-random structure within these branching patterns such that neurons with four targets are more frequently observed than those with two or three, indicative of widespread networks. Consequently, these multi-target single neurons form distinct networks within medial and ventral frontal cortex consistent with their known functions in regulating mood and decision-making. Additionally, we show that branching patterns of single neurons shape functional networks in the brain as assessed by fMRI-based functional connectivity. These results provide a neuroanatomical basis for the role of the BLA in coordinating brain-wide responses to valent stimuli^8^ and highlight the importance of high-resolution neuroanatomical data for understanding functional networks in the brain.

## Introduction

Basolateral amygdala (BLA) is essential for adaptive emotional responding in humans and animals^5, 9, 10^. In humans, dysfunction within or damage to the circuits that connect through the amygdala are theorized to be the cause of numerous psychiatric disorders, including autism spectrum disorder, post-traumatic stress disorder, and schizophrenia^11–13^. Based on decades of tract-tracing studies in macaques, we know that the primate BLA projects widely across the brain, sending connections primarily to ventral and medial parts of the frontal lobe, as well as the temporal and occipital cortices, thalamus, and striatum^1–4^. These diverse and widespread connections, especially those to the frontal lobe, are central to accounts of how BLA in humans coordinates learning about and responding to different emotionally salient events^14, 15^.

Despite the appreciation that the BLA plays a central role in coordinating activity across large networks to guide emotional behavior, the anatomical organization of single neuron connections from this area are largely unknown. One possibility is that single BLA neurons project to only one specific target, transmitting information to downstream targets in dedicated pathways. Such an organizing principle or connectional motif would align closely with the BLA’s known role in model-based behaviors and the processing of sensory-specific stimuli through interaction with distinct parts of frontal cortex^16, 17^. An alternative is that single neurons in BLA branch to many different areas, such that activity can be efficiently coordinated across distributed networks of areas. This organization fits with the BLA’s role in more general aspects of motivation and response invigoration to approach or avoid salient stimuli that are characteristic of model-free behaviors^8^.

These two potential connectional motifs of single BLA neurons – specific vs branching – are not necessarily mutually exclusive. However, at present, the extent to which either motif best characterizes the projections of single BLA neurons or indeed populations of BLA neurons is not known, in part because gold-standard tract-tracing approaches are either too coarse to detect the projection patterns of individual neurons^18, 19^ or because the available single-axon tracing techniques do not scale practically to non-human primates^20^. To surmount these issues, we optimized and refined a high-throughput sequencing approach, multiplexed analysis of projections by sequencing (MAPseq^7^) in macaque monkeys. MAPseq uses barcoded mRNA technology^21^ to map the connections of individual neurons at scale. Because of their potential importance in psychiatric disorders^11^, we focused on projections from BLA to frontal cortex, striatum, anterior temporal lobe, and mediodorsal nucleus of the thalamus (MD), a part of thalamus that receives input from both frontal cortex and amygdala^1^. Using this approach, we found that individual BLA neurons project widely in frontal cortex; about half of the neurons that leave amygdala project to more than one target area. The pattern of these branching projections was not random, such that the connections of single neurons were organized into distinct and reproducible connection motifs. Notably, BLA projections to posterior parts of frontal cortex were highly specific, whereas those to more anterior parts of frontal cortex, especially ventral frontal cortex, were more likely to branch to multiple areas.

## Results

### Optimization of MAPseq in macaques

MAPseq^7^ relies on an engineered sindbis virus that infects neurons with unique RNA sequences, referred to as barcodes^21^. Following viral expression these barcodes are conjugated to a nonfunctional presynaptic protein and undergo anterograde axonal transport. Thus, by dissecting and sequencing samples from injection and target brain areas, the projection patterns of single neurons can be determined. MAPseq thereby complements and extends traditional neuroanatomical approaches, as it provides both information about bulk projection patterns from one area to another as well as the connection patterns of single neurons. Further, it permits simultaneous analysis of projections to many target areas within the same animal, allowing single-neuron branching to be discerned.

We performed bilateral, MRI-guided stereotactic injections of barcoded sindbis virus into the BLA of two rhesus macaques (**Figure 1A**, see Extended Data Figure 1 for more detail); 10 to 12 injections of 400 nl each were placed throughout the lateral, basal, and accessory basal nuclei to control viral spread. Following perfusion, brains were extracted, the hemispheres separated and sectioned. The BLA and target areas in frontal cortex, striatum, entorhinal cortex, hippocampus, and MD were dissected according to gray/white matter boundaries as well as sulcal landmarks (**Figure 1B**). Simple qPCR on extracted mRNA recovered significantly more barcode in amygdala sites near the injection compared to target sites (OLS regression, t(2) = 46.43, *p* < 0.0001) and more in target sites compared to control sites in cerebellum (t(2) = 4.77, p < 0.0001; **Figure 1C**). High-throughput next-generation sequencing was then conducted on extracted mRNA from BLA and target areas. Counts of unique barcodes from the extracted RNA were normalized and a threshold applied to control for spurious sequencing results^7^ (Extended Data Figure 2). The combined thresholded barcode counts from the four hemispheres were then analyzed together (see Extended Data Figures 3 and 4 for analyses of each hemisphere separately and comparisons between hemispheres, respectively).

**Figure 1:**
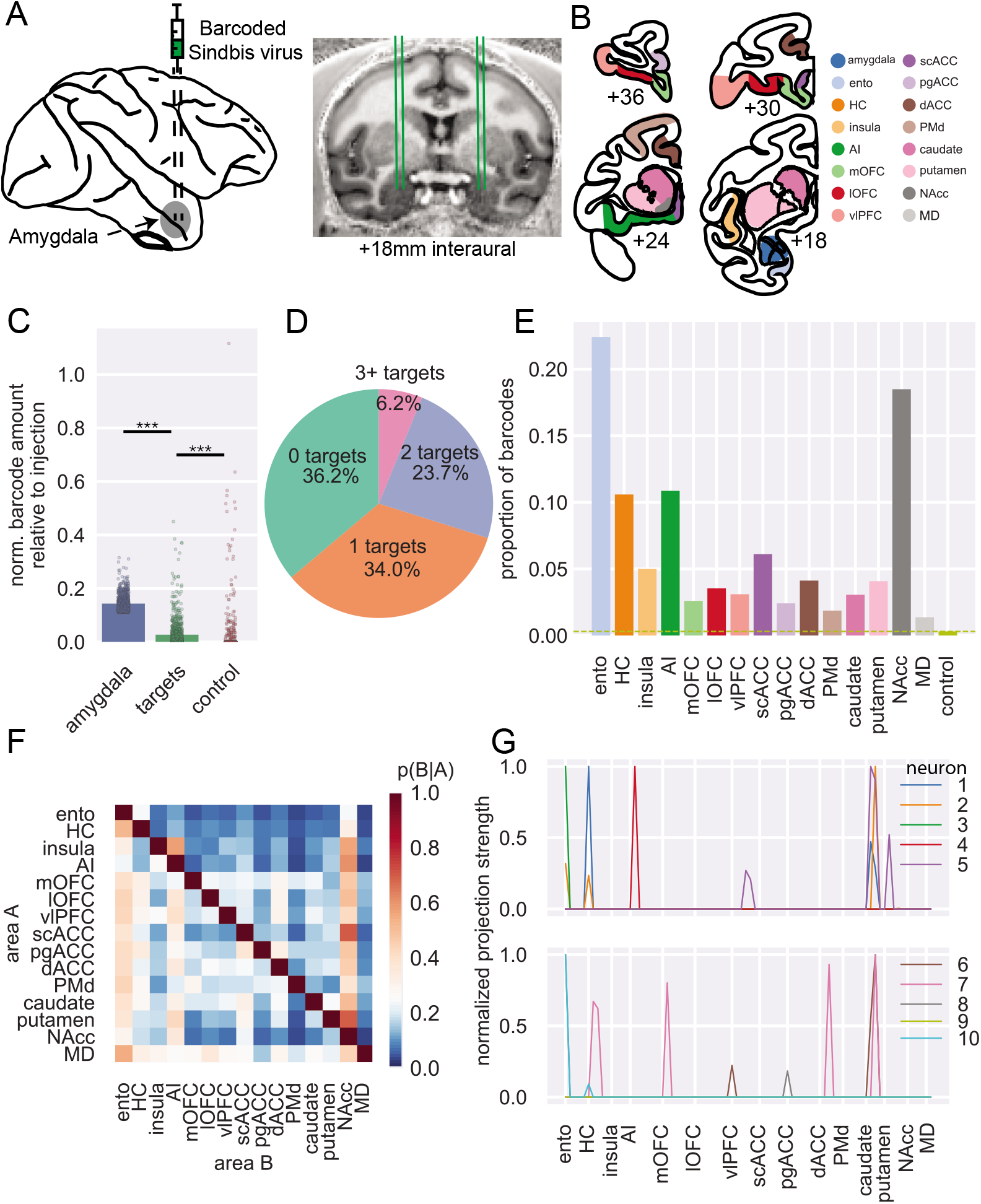
MAPseq of macaque BLA. **A**) Schematic of injection approach. **B**) Example coronal sections showing dissection targets in striatum as well as frontal and temporal lobes. Ento, entorhinal; HC, hippocampus (not shown); AI, anterior insula cortex; mOFC, medial orbitofrontal cortex; lOFC, lateral orbitofrontal cortex; vlPFC, ventrolateral prefrontal cortex; scACC, subcallosal anterior cingulate cortex; pgACC, perigenual ACC; dACC, dorsal ACC; PMd, dorsal premotor cortex; NAcc, nucleus accumbens; MD, mediodorsal thalamus (also not shown). Anterior-posterior levels are in mm relative to the interaural plane. **C**) Normalized barcode amount relative to peak barcode amount in injection site across other locations in amygdala (blue), all target structures (green; OLS regression vs amygdala: t(2) = 46.43, *p* < 0.0001), and control areas (red; vs targets: t(2) = 4.77, *p* < 0.001). **D**) Proportion of unique barcodes in no targets outside of amygdala (green), one (orange), two (blue), or more than two targets outside of amygdala (red). **E**) Proportion of barcodes in each of the target and control areas; dashed line indicates the proportion of barcode found in control sites. **F**) Conditional probability that a barcode in area A is also found in area B. Note that the order of indexing here alters the probability such that P(A|B) can be greater than P(B|A). **G**) Normalized proportion of barcodes across all the potential target areas and control areas for 10 unique barcodes (colored lines).

In total, we recovered 3,115 unique barcodes in samples dissected from BLA across the four hemispheres. This yield of barcode counts is similar in level to that recovered in mice per injection^21–24^, indicating that MAPseq^7^ was working as intended in macaques. To determine the projection targets of putative single neurons from BLA, we then determined whether each barcode found in amygdala was present in any of the target areas after collapsing across all samples in a target area. Approximately one-third of the unique barcodes – which can be interpreted to represent single neurons – projected only to other sites within amygdala, which we refer to as zero-target. Another third projected to only one of the target areas we collected – we refer to these neurons as having *specific* projections – and the remaining third had two or more targets outside of amygdala. We refer to these neurons with multiple targets as having *branching* projections (**Figure 1D**).

The overall proportion of barcode recovered in each target mirrored known connections of BLA in macaques (**Figure 1E**). Accordingly, the highest amount of barcode was recovered in entorhinal cortex and nucleus accumbens (NAcc), two areas which have previously been identified as receiving dense projections from BLA^25^. A lower amount of barcode was recovered from hippocampus and agranular insula (AI), which are also well-documented as receiving strong projections from BLA^4, 26^, followed by subcallosal (scACC) and dorsal ACC (dACC). Notably, relatively similar amounts of barcode were recovered from all other target areas in frontal cortex and striatum, which again matches known gradients of BLA connections across the frontal lobe^4, 27^. The close alignment between our findings and prior tract-tracing supports the validity of MAPseq in quantifying coarse area-to-area projections in macaques.

Next, we looked at the probability that a barcode found in one target structure was also found in another target structure (**Figure 1F**). This approach allowed us to begin to ascertain the degree to which connections of single BLA neurons are either specific or branching. For instance, neurons that project to NAcc have a high probability of also projecting to AI. By contrast, single neurons that project to AI are unlikely to project to either entorhinal cortex or hippocampus. Similar patterns of non-overlapping projections are also apparent in BLA projections to frontal cortex; neurons that project to medial orbitofrontal cortex (mOFC) are unlikely to project to ventrolateral prefrontal cortex (vlPFC). As can be seen in by the representative examples in **Figure 1G**, evidence for both specific and branching projection patterns can be observed at the level of single neurons, as well. Notably, our results correspond closely with the findings of two studies that investigated the patterns of BLA neuron branching to frontal cortex and thalamus. One by Sharma, Fudge and colleagues^28^ identified a subset of amygdala neurons that project to adjacent parts of medial frontal areas 25/14 (which we refer to as scACC and mOFC, respectively) and 32/24 (which encompasses our perigenual ACC [pgACC] and dACC, respectively). We also identified neurons that projected to both areas, although we identified a slightly higher proportion of neurons with this branching pattern (between 7 and 20% depending on the animal) of them than previously reported (between 7 and 10% depending on the nucleus) – likely because our dissected areas encompassed larger portions of medial frontal cortex. A study by Timbie and Barbas^29^ found that two largely non-overlapping populations of amygdala neurons projected to either MD or posterior OFC/AI, a result highly similar to what we found (**Figure 1F**). This lack of branching as discerned by our barcode analysis is an important negative finding; in additional to observing the expected pattern of *branching projections* to certain targets, we have also observed the expected pattern of *specific projections* to other targets. Taken together the above analyses provide novel evidence that BLA neurons often project to multiple targets in frontal cortex, arguing against specific targeting as the dominant connectivity principle, while also reproducing the known gross connectivity of BLA – altogether confirming the validity of MAPseq in macaques.

### Quantitative analysis of branching connectional motifs of single BLA neurons

The prior analysis on the branching of neurons to two targets, while revealing, does not capture the full set of connections that single neurons make to multiple target areas – one of the major strengths of MAPseq over standard tract-tracing approaches (**Figure 1G**). Consequently, we focused our next analysis on the nearly 1,300 BLA neurons with branching projections to multiple locations in frontal cortex, temporal cortex, striatum and thalamus (**Figure 2A**). While all target areas received the majority of their input from branching neurons, entorhinal cortex was found to receive the highest proportion of specific input from BLA (**Figure 2B**, z-test for proportions: entorhinal vs hippocampus [next highest], z = 10.67, *p* < 0.0001). Next, to determine whether any branching motifs were over- or under-represented compared to chance, we built a null distribution based on the overall barcode distribution (**Figure 2C**). Assuming total independence for branching, the probability that a neuron projects to both areas A and B can be computed as the product of the independent probabilities of projecting to area A and area B^24^. By comparing the actual counts for each motif with this null distribution, we identified only one over-represented bifurcating motif: neurons that project to both NAcc and AI (binomial test, *p* = 0.003). This motif was found alongside predominantly under- represented bi- and trifurcations, many of which included projection motifs encompassing some combination of NAcc, entorhinal cortex, and hippocampus (**Figure 2D and 2E**). That branching to only two of these areas happens less frequently than expected is somewhat surprising considering these were the three areas most likely to receive projections from BLA in our data (**Figure 1E**); however, these data are in line with small-world theories of brain network organization^30^. By contrast, axon branching to four different targets were far more likely to be over-represented compared to that same chance distribution (**Figure 2F**). In other words, there are fewer bifurcations than expected based solely on the proportions of barcodes found in each target area, while there were more of these quadfurcations than expected. There were no significantly over- or under-represented motifs with five or more targets. Thus, these results indicate that single BLA neurons demonstrate a high degree of branching, being more likely to strongly innervate four distinct targets in the target areas sampled over two or three. Importantly, this analysis also reveals that the observed branching motifs are not simply a product of the distribution of barcodes; rather, branching of single BLA neurons appears to be highly structured.

**Figure 2:**
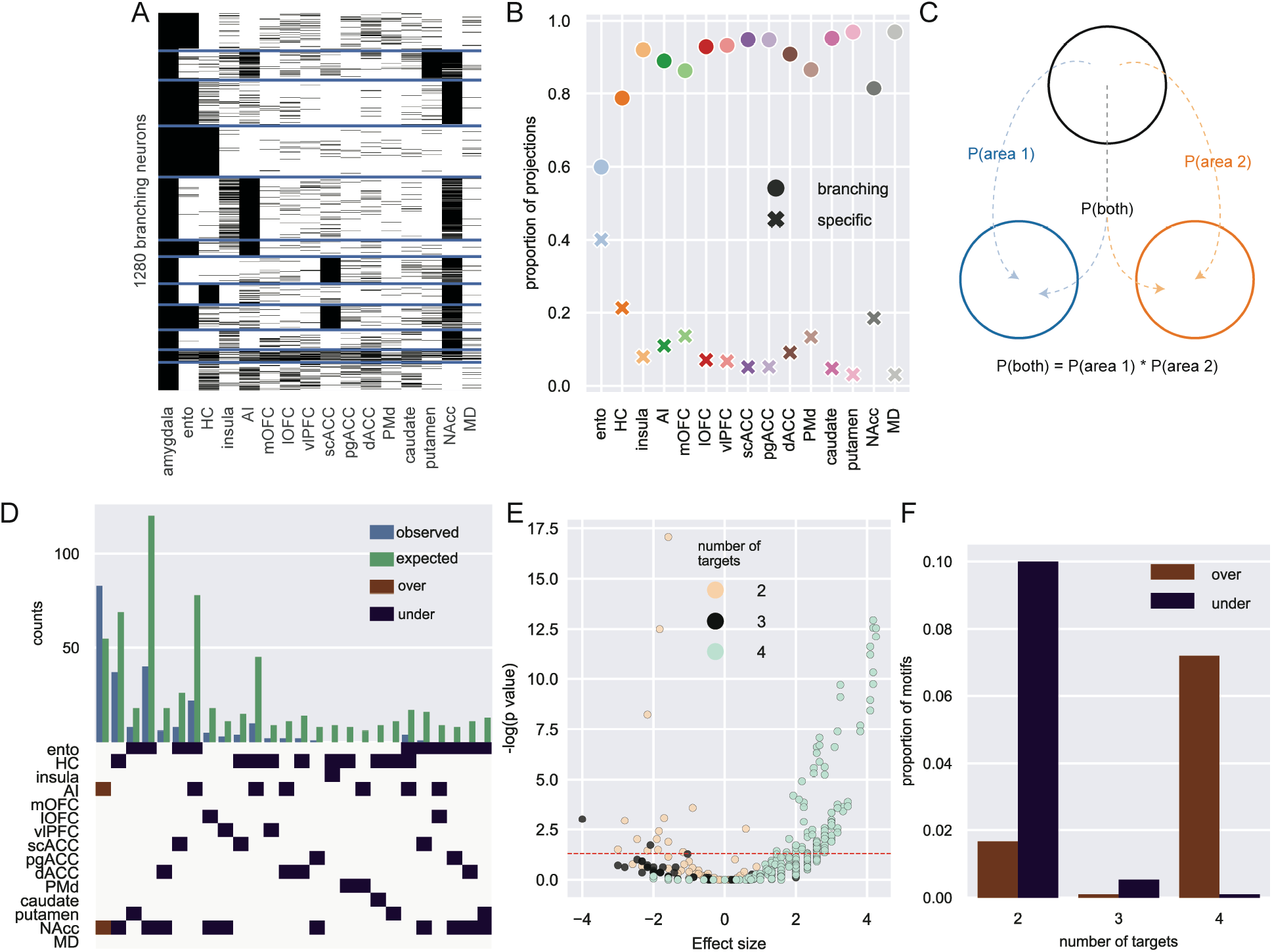
Single neuron analysis of branching projections from BLA. **A**) K-means clustering of 1,300 neurons that project to more than one area outside of amygdala (k=12). **B**) Proportion of projections from BLA to frontal cortex areas, striatum, and mediodorsal thalamus that are from specific (crosses) or branching projections (filled circles). **C**) Logic of null distribution computation for branching motifs. **D**) Observed (blue) and expected (green) counts of neurons with projections to multiple areas (top). Specific over (red) or under-represented (blue) branching motifs by area (bottom). **E**) Volcano plot of probability of all possible branching motifs with 2 (cream), 3 (grey), and 4 (turquoise) target areas. Dashed line marks the level of statistical significance. **F**) Proportion of significantly over- (red) and under- (blue) represented 2-, 3- and 4- target area branching motifs.

### Single neuron projection networks within frontal cortex

With the appreciation that the projections of single neurons in BLA are highly likely to branch to multiple areas we sought to understand how these projections are organized. Here we separately focused on the pattens of single BLA projections to the medial and ventral frontal cortex. We took this approach because BLA projections to these areas are thought to be functionally distinct. Interaction between BLA and medial frontal cortex is heavily linked to defensive threat conditioning in animals and anxiety-related disorders such as PTSD in humans^31^. Projections from BLA to ventral frontal cortex are, by contrast, more frequently associated with reward-guided behaviors^6^ and dysfunction in these circuits is linked to addictive disorders^32^.

#### Medial frontal cortex

We identified 405 BLA neurons that project to either medial frontal areas scACC (area 25 as defined by Carmichael and Price^33^), pgACC (area 32), or dACC (area 24) (**Figure 3A**). Although these areas are densely interconnected^18^, we found marked differences in the structure of BLA input that they receive. First, over half of the BLA neurons that projected to medial frontal cortex exclusively targeted scACC, whereas a third targeted only dACC (130/405; z-test for proportions, z = 6.38, *p* < 0.0001); even fewer BLA projections to medial frontal cortex were specific to pgACC (55/405; z = 6.28, *p* < 0.0 001) (**Figure 3B**). Indeed, the majority of BLA neurons targeting pgACC also projected to the other parts of medial frontal cortex, while a smaller proportion of dACC-projecting neurons branched within medial frontal cortex (z = 5.37, *p* < 0.0001); scACC-projecting neurons were least likely to branch (z = 3.17, *p* = 0.0015). Thus, within medial frontal cortex there is a hierarchy of specific vs branching BLA connections where pgACC receives the least specific BLA input and scACC receives the most.

**Figure 3:**
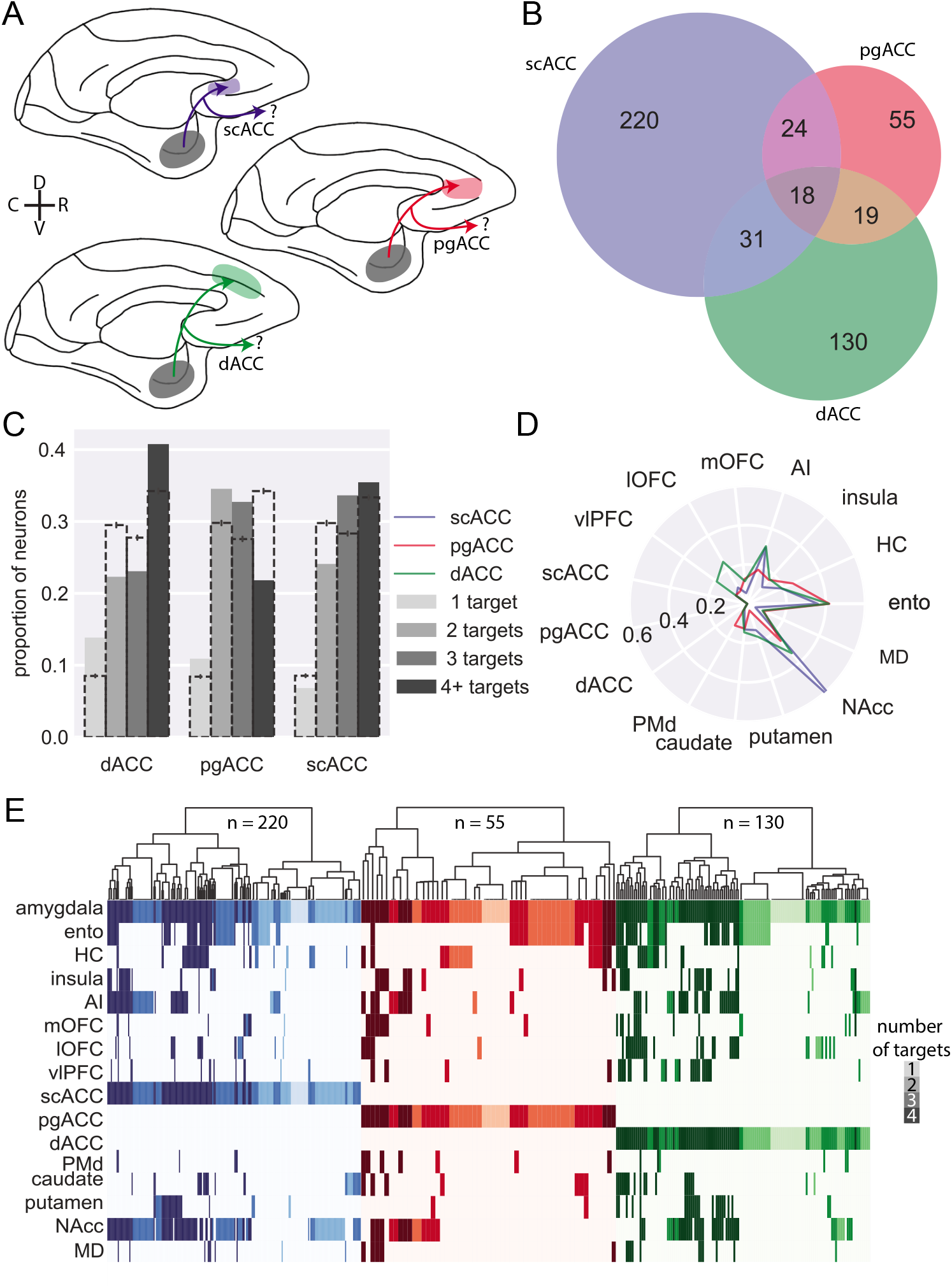
BLA specific projections to medial frontal cortex. **A**) Schematic showing the three populations of medial frontal cortex-projecting neurons analyzed here: BLA neurons targeting scACC (blue), pgACC (red), dACC (green). D and V refer to the dorsal and ventral directions, respectively; R and C designate rostral and caudal. **B**) Venn diagram illustrating projections to each medial frontal cortex area that also branch to the other two; pgACC- projecting neurons are more likely to branch than dACC-projecting neurons (z-test for proportions, z=3.17, p=0.0015), which are more likely to branch than scACC-projecting neurons (z=2.27, p=0.023). **C**) Degrees of branching for each medial frontal cortex-projecting population; dashed bars indicate the mean of 1000 shuffles of the data, downsampled for equal numbers of neurons from each population; error bars indicate 95% confidence intervals. **D**) Likelihood of medial frontal cortex-projecting neurons projecting to non-medial frontal cortex targets. **E**) Single-neuron projection patterns; shading indicates number of targets.

As we noted earlier, BLA neurons tend to have more branching than specific projections (**Figure 2C** and **E**); within this branching, we found additional structure among projections that targeted single medial frontal areas. Neurons that project to scACC or dACC were more likely to have four targets than neurons which projected to pgACC, which were dominated by two- and three- target neurons (**Figure 3C**, permutation test). These results suggest that amygdala inputs to pgACC, while frequently shared among other cingulate areas, are not shared as frequently outside of medial frontal cortex.

Within the projections from BLA to medial frontal cortex there were notable differences in the areas that these single neurons also targeted, indicative of different networks. Whereas the BLA neurons projecting to each medial area had similar proportions of bifurcations to hippocampus and entorhinal cortex (**Figure 3D** and **E**), neurons that projected to scACC were far more likely to also project to NAcc than the other areas (z-test for proportions: scACC vs pgACC: z = 5.85, *p* < 0.0001; scACC vs dACC: z = 6.33, *p* < 0.0001). This projection motif is evident at the level of individual neurons, and these cells were also highly likely to connect to AI (**Figure 3E**). dACC-projecting BLA neurons, however, were more likely to also project to lateral OFC (lOFC; dACC vs pgACC: z = 2.72, *p* = 0.0097; dACC vs scACC: z = 3.80, *p* < 0.001) and vlPFC (dACC vs pgACC: z = 2.04, *p* = 0.061; dACC vs scACC: z = 3.52, *p* < 0.001) on the ventral surface of the frontal lobe, providing further anatomical support for the role of the dACC in valuation through its interactions with more ventral areas^6, 34^. In summary, single BLA neurons targeting medial frontal areas appeared to target distinct networks; those targeting scACC were largely constrained to this area and primarily sent bifurcations to NAcc and AI in the posterior ventral frontal cortex, consistent with the known roles of these areas in regulating mood^35^. By contrast, those targeting dACC also innervated parts of ventral frontal cortex including those linked to Carmichael and Price’s visceromotor network^18^, consistent with its role in value-based decision- making^6, 34^.

### Ventral Frontal Cortex Networks

We conducted a similar analysis on the 627 BLA neurons that projected to areas on the ventral surface of frontal cortex, including AI, mOFC, lOFC, and vlPFC (**Figure 4A**). Of those BLA neurons that projected to ventral frontal cortex, almost two thirds (65%) of BLA neurons solely targeted AI, whereas only 10% solely targeted vlPFC; within ventral frontal cortex, these areas received the highest and lowest proportions of specific projections, respectively (z-test for proportions: z = 19.85, *p* < 0.0001, **Figure 4B**). Indeed, the majority of BLA neurons projecting to vlPFC also targeted other parts of the ventral frontal cortex and were not specific to this area. BLA neurons projecting to mOFC and lOFC had similar proportions of cells projecting in a specific or branching manner. Thus, BLA projections to AI are more specific compared to those directed to more anterior regions (AI vs mOFC: z = 4.43, *p* < 0.0001), with vlPFC receiving the fewest specific projections from BLA (vlPFC vs mOFC: z = 2.48, *p* = 0.013).

**Figure 4:**
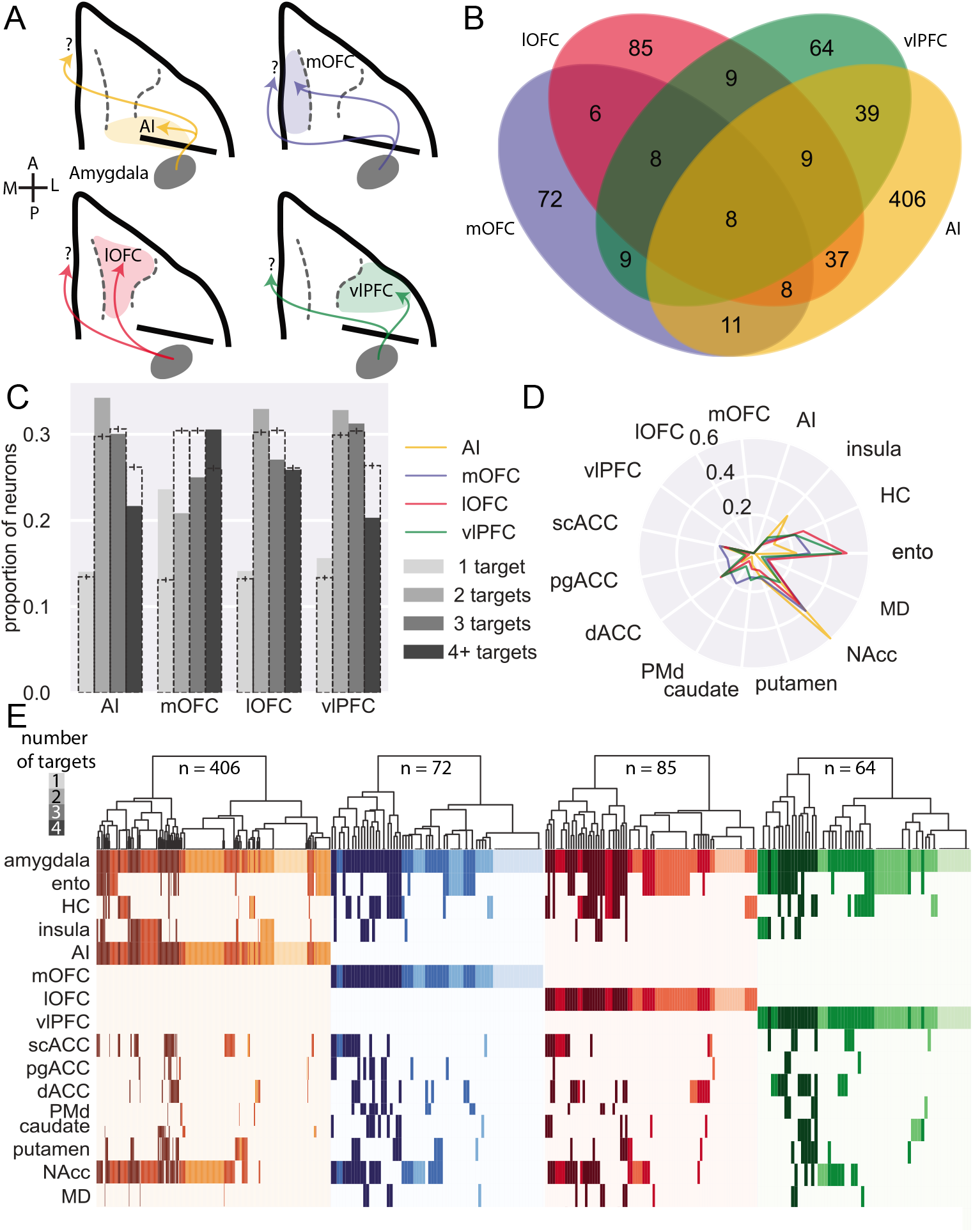
BLA specific projections to ventral frontal cortex. **A**) Schematic showing the four populations of ventral frontal cortex-projecting neurons analyzed here: BLA neurons targeting AI (yellow), mOFC (blue), lOFC (red), and vlPFC (green). A and P designate anterior and posterior directions, respectively; M and L refer to medial and lateral. **B**) Venn diagram illustrating projections to each area that also branch to the other three; AI-projecting neurons were least likely to branch to the other areas (z-test for proportions, z=4.425, p<0.0001), while vlPFC- projecting neurons were most likely to branch within ventral FC (z=2.476, p=0.0133). **C**) Degrees of branching for each ventral frontal cortex-projecting population; dashed bars indicate the mean of 1000 shuffles of the data, downsampled for equal numbers of neurons from each population; error bars indicate 95% confidence intervals. **D**) Likelihood of ventral frontal cortex-projecting neurons projecting to non-ventral frontal cortex targets. **E**) Single-neuron projection patterns; shading indicates number of targets.

Similar to medial frontal cortex, BLA neurons projecting to ventral frontal cortex were more likely to send branching as opposed to specific projections. AI-, lOFC-, and vlPFC- projecting neurons were most likely to send 2-target projections (**Figure 4C**, permutation test), whereas mOFC-projecting neurons tended to innervate either one or four targets. AI- and vlPFC-projecting neurons were least likely to branch to four distinct targets, suggesting that these two areas receive the most highly specialized input from BLA.

When we limited our analyses to BLA neurons that only projected to one of the four ventral frontal areas without branching between them, BLA neurons that project to AI exhibited the strongest projections to NAcc compared to other areas in ventral frontal cortex (**Figure 4C** and **E;** Fisher’s exact test: AI vs mOFC, *p* = 0.0057; AI vs lOFC, *p* = 0.00019; AI vs vlPFC, *p* < 0.0001). By contrast, vlPFC- and lOFC- projecting neurons were most likely to also project to entorhinal cortex compared to mOFC and AI (**Figure 4C;** lOFC vs mOFC, *p* = 0.043; lOFC vs AI, *p* < 0.0001; vlPFC vs AI, *p* = 0.00051). Somewhat unexpectedly, BLA neurons with projections to mOFC were more likely to project to dorsal premotor cortex (PMd; mOFC vs AI, *p* < 0.00001; mOFC vs lOFC, *p* = 0.028) and other more medial areas of the frontal lobe such as the pgACC (**Figure 4C** and **E;** mOFC vs AI, *p* = 0.019; mOFC vs lOFC, *p* = 0.038) than were BLA neurons projecting to other ventral frontal cortex regions. These patterns of projections indicate the BLA neurons targeting the ventral frontal cortex form highly-structured networks; those targeting AI are less likely also to project to other ventral frontal areas. When they do branch to other areas, they primarily innervate other areas in the posterior frontal lobe (i.e. scACC) and NAcc. By contrast, BLA projections to more anterior parts of ventral frontal cortex, especially vlPFC, branch more both within frontal cortex and also to other areas. This finding that vlPFC- projecting neurons connect so broadly aligns closely with the known role of the BLA-vlPFC circuit in representing and updating model-free stimulus-outcome associations^36^, information that is likely to be shared among other ventral frontal areas to inform reward-guided choice behavior^37^.

Taken together, the patterns of projections strongly indicate the existence of BLA neurons that target distinct networks in the medial and ventral frontal cortex. Those BLA neurons that target scACC and AI in the posterior frontal cortex preferentially innervate NAcc. By contrast, BLA neurons that target to more anterior medial and ventral frontal areas appear to be part of more distributed networks of areas. This extends our understanding of the organization of BLA-to-frontal cortex networks revealing distinct connectional motifs that may in part explain the pervasive influence of BLA on frontal cortex processing.

### Retrograde tracing validation of specific and branching BLA projections to ventral frontal cortex

Next, we sought to assess the most distinct BLA projection patterns revealed by MAPseq using standard retrograde viral tracing. In a single macaque, MRI-guided stereotactic injections of retro-AAV2 coding for mCherry and EGFP fluorophores were injected into NAcc and a lateral subregion of AI in one hemisphere. Injections were targeted to posterior-lateral lOFC (area 13m as defined by Carmichael and Price^33^) and posterior-medial vlPFC (area 12o) in the other hemisphere (**Extended Data Figure 5A and B**). As cross-hemispheric BLA connections are negligible^27^, we were able to analyze the two hemispheres separately. We then conducted unbiased stereological counting of neurons in BLA that were either single or double labeled with each fluorophore in each hemisphere^38^. Thus, we were able to compare the connectivity profiles of single amygdala neurons found in specific subregions of frontal cortex.

For BLA neurons projecting to AI, we found a high degree of correspondence in the proportion of branching neurons between stereology and MAPseq estimates (z-test for proportions *p* > 0.05, Extended Data Figure 5C); for NAcc-projecting neurons, however, our MAPseq and stereology results did not agree, as stereological estimates of specific projections were higher than MAPseq (*p* < 0.0001). For lOFC and vlPFC, too, the numbers of BLA neurons that projected specifically to one or branched to both areas was different to what we found with MAPseq (*p* < 0.001, Extended Data Figure 5D). This was primarily because stereological estimates of BLA projections to vlPFC almost entirely overlapped with those targeting lOFC. The difference in estimates of specific versus branching BLA projections between retrograde tracing and MAPseq results is not unexpected^22^. This is because the injections of retro-AAV into lOFC and vlPFC do not cover the full extent of these cortical regions. By contrast, MAPseq estimates of specific and branching projections are based on BLA projections to the whole of lOFC and vlPFC that extend far beyond the extent of the areas target with retro-AAV. The difference in estimates of the branching and specific projections between techniques is, however, revealing as it indicates that projections to this subregion of area 12 are even less specific than the analysis of connectivity of the larger vlPFC would indicate.

### Branching and specific projections shape fMRI functional connectivity

Anatomical connections in the brain constrain functional networks^39, 40^. Given the reproducible patterns of specific and branching projections from BLA that we identified, we sought to determine if multi-area connectional motifs identified by MAPseq could be identified at the level of fMRI functional connectivity (FC). If such patterns were identifiable in FC this would indicate that the unique anatomical networks identified with MAPseq are related to and in fact shape functional networks in the brain. We analyzed a dataset of resting state fMRI scans from six rhesus monkeys^41^ (**Figure 5A**), with single seed voxels in amygdala and target voxels in the same areas used in the MAPseq experiments (Extended Data Figure 6). After computing and z-transforming FC values for each amygdala voxel with each target voxel, we binarized the FC signal by setting a threshold at 70% of each voxels’ maximum connectivity. Practically, this meant that if FC between an amygdala voxel and any target area was above the threshold, we counted that amygdala voxel as being *functionally connected* to that target area. We then counted the number of connections to each target area and normalized those counts by the size of each target area. This approach meant that single amygdala voxels could be designated as being functionally connected to multiple parts of frontal cortex, striatum, temporal lobe and thalamus, following a similar approach to MAPseq.

**Figure 5:**
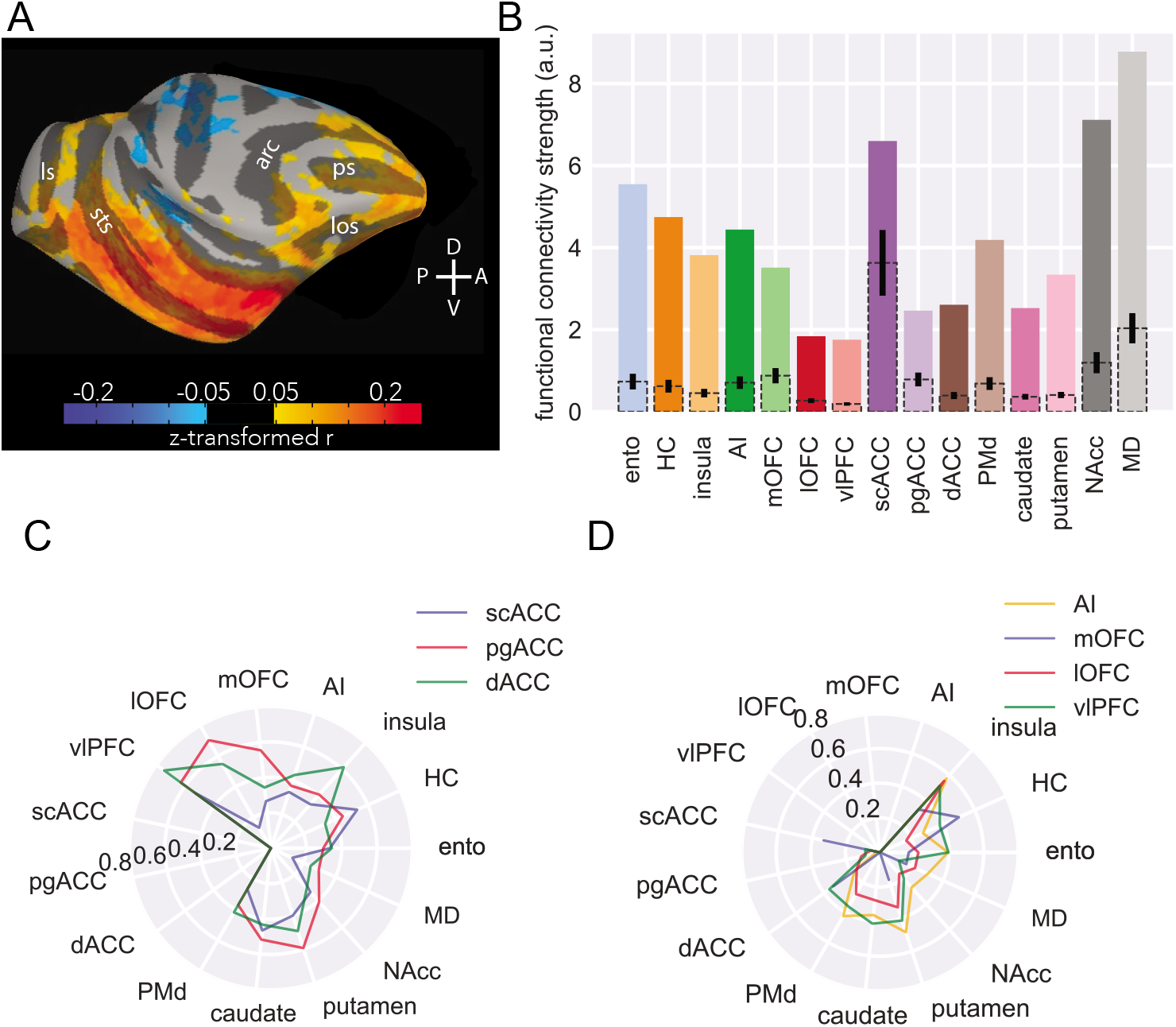
MRI functional connectivity replicates some anatomical features. **A**) Surface projection of right amygdala ROI functional connectivity to right hemisphere targets, averaged across six animals. Compass directions refer to dorsal (D)/ventral (V) and posterior (P)/anterior (A) directions; sulci labelled are: principal sulcus (ps), lateral orbital sulcus (los), arcuate sulcus (arc), superior temporal sulcus (sts), lunate sulcus (ls). **B**) Functional connectivity strength across areas. Colored bars indicate number of amygdala voxels functionally connected to each target area, corrected for the number of voxels in each target area; dotted bars with 95% confidence intervals indicate random samples of the same number of voxels from each area as the smallest target area (scACC). **C**) Likelihood of medial FC-projecting voxels being functionally connected to non-medial FC targets. **D**) Likelihood of ventral FC-projecting voxels being functionally connected to non-ventral FC targets.

Despite fMRI-derived FC operating over a much larger spatial scale than standard anatomical approaches and having no directional sensitivity, we were still able to identify a largely similar pattern of relative FC between amygdala and our target areas as we had found with MAPseq (compare **Figures 1E** and **5C**). Notably, the pattern also generally matches prior tract-tracing results as there was strong connectivity between amygdala and entorhinal cortex, hippocampus, NAcc, AI, scACC as well as MD with lower connectivity to lOFC and pgACC^42^. For a number of areas, functional connectivity with amygdala was higher than expected based on MAPseq and tract-tracing data, most notably PMd and mOFC (**Figure 5C**). This result is not necessarily surprising because FC measures not only reflect unidirectional connections from one area to another; rather, strong functional connection between two areas can be driven by strong reciprocal connections or strong coupling with a third area with which both areas strongly interact^43^.

Next to look for patterns of network-level connectivity we assessed the FC of amygdala voxels with high connectivity to areas in medial frontal cortex (**Fig. 5C**) or ventral frontal cortex (**Fig. 5D**). This is essentially the same analysis depicted in **Figures 3E** and **4E**. Notably, we were able to identify many of the prominent connectivity motifs that we observed with MAPseq. For instance, voxels with a high likelihood of being connected to AI neurons also had higher connectivity with NAcc (**Figure 5D**). Similarly, dACC-connected neurons also had a high likelihood of being connected to vlPFC (**Figure 5C**) and if a voxel showed high FC with any of the medial and ventral frontal cortex areas it had uniformly low connectivity with MD (**Figures 5C** and **D**). This latter point demonstrates the correspondence between MAPseq, tract-tracing and fMRI^42^.

Despite being able to discern these broad connectional motifs, some key differences between medial and ventral areas were obscured, such that fMRI estimates of multi-region connectivity estimated highly similar connectivity profiles for all three medial frontal cortex areas. This difference between fMRI and MAPseq analyses is likely explained by the dense and bidirectional interconnections within medial frontal cortex^18^, inputs from other structures received by this part of frontal cortex most notably from hippocampus and neuromodulatory systems^27, 44^, and the low resolution of fMRI compared to individual neurons. Taken together, that major anatomical features identified by MAPseq were also identifiable at the level of fMRI FC likely reflects their significance in shaping brain-wide activity patterns organized into functional networks.

## Discussion

We have optimized and validated MAPseq for single-neuron connectivity mapping in macaque monkeys, opening new avenues of research into the single-cell structural connections of the non-human primate brain. Using this approach, we successfully determined the projection patterns of over 3,000 single neurons from BLA to frontal cortex, striatum, parts of the temporal lobe, and MD. Notably, the bulk patterns of connections identified here are in close alignment with previous reports of BLA projections based on traditional tract-tracing techniques^1,2,3, 25, 26, 29, 45^ (**Figures 1 and 2**). Overall, we found that single BLA neurons branch extensively, with branching to four distinct targets being more likely to be over-represented compared to chance than branching to two or three targets (**Figure 2**). Within these patterns of branching, we identified distinct connectional motifs; BLA projections to posterior parts of the medial and ventral frontal cortex were highly specific and less likely to branch to other areas. By contrast, projections to more anterior or dorsal areas were much more likely to branch to other brain areas and exhibited unique projection profiles (**Figures 3 and 4**). We also identified broad similarities and slight differences between this anatomical connectivity and more standard tract-tracing techniques (**Extended Data Figure 5**) and functional connectivity identified by fMRI (**Figure 5**). These findings begin to reveal the unique patterns of projections of single BLA neurons, connections that are heavily implicated in the control of affect and that become dysfunctional in psychiatric disorders^11^.

It is well documented that BLA input to frontal cortex is strongest to posterior regions and weaker to more anterior regions^2, 4^. On top of this, we found that in both medial and ventral frontal cortex, BLA neuron branching is highest to the most anterior areas. For instance, BLA neurons projecting to pgACC showed the highest degree of branching to other areas in medial frontal cortex, whereas neurons projecting to the more posterior scACC branched the least (**Figure 3B**). A generally similar pattern was seen in ventral frontal cortex with the exception of vlPFC, which received the highest degree of branching projections (**Figure 4**). This lack of specific BLA projections to vlPFC may be related to its role in model-free as opposed to model-based behaviors; lesions of vlPFC do not impact reinforcer devaluation that depends on specific sensory information about an outcome but do impact outcome-independent probabilistic learning^36^. Thus, the patterns of specific and branching connections from single BLA neurons may serve a functional role in relaying distinct sensory information or providing a salience signal to invigorate responding, respectively. In medial frontal cortex, while so-called ventromedial PFC has been generally implicated in affective regulation, and by extension anxiety and mood disorders^46^, more recent evidence suggests that particular subregions may play distinct roles in affect^47–49^. The highly specific and segregated inputs to scACC and pgACC identified here potentially provide a neuroanatomical basis for their opposing roles in affective responding^35^.

Appreciating the unique features of BLA projections to frontal cortex is potentially critical for understanding the basis of a number of amygdala-linked psychiatric disorders^11^. For example, obsessive-compulsive disorder is associated with dysfunction in basal ganglia-OFC circuitry regulating valuation as well as basal ganglia-dACC circuits involved in action selection^50^. Here we identified a population of dACC-projecting BLA neurons that also preferentially targeted parts of ventral frontal cortex including lOFC (**Figure 3**). Our findings therefore provide a potential anatomical basis through which dysfunction in a single, small population of BLA neurons could influence a distributed network of areas. Determining the distinct functions of the different BLA projection motifs identified here as well as their molecular signatures^23, 51^ has the potential to bring network-level understanding to basic and translational neuroscience and might provide a more biologically-realistic basis for the construction of neural network architectures^52^.

## Supporting information

Supplemental figures only

## Author Contributions

ZRZ, RLC, PRH, FMS, and PHR conceived the project. ZRZ and PHR designed and carried out the experiments, performed the analyses, and wrote the paper. LL performed the stereology analysis, WGJ assisted in tissue preparation, JMF assisted in surgery, CE and AF collected the fMRI scans, HZ performed the RNA extraction and sequencing, and BER performed the fMRI preprocessing and analysis. RLC, PRH, and FMS provided vital feedback on analysis approaches throughout. All authors approved the final version of the paper.

## Acknowledgements

ZRZ, RLC, PRH, and PHR are supported by a grant from the BRAIN initiative (R34NS122050). We thank Allison Sowa of the ISMMS Microscopy and Advanced Bioimaging CoRE for help with tissue preparation, members of the Rudebeck Lab for assistance with surgical procedures, and the MAPseq Core at CSHL for assistance in tissue processing and sequencing. We also thank Anthony Zador and Alex Vaughan for advice and encouragement and Elisabeth Murray for comments on an earlier version of the manuscript.

## Methods

### Subjects

Two male rhesus monkeys (*Macaca mulatta*) and one male long-tailed macaque (*Macaca fascicularis*), 8-9 years of age, were used for our experiments, weighing between 10 and 15 kg. The two rhesus macaques were used in MAPseq experiments whereas the long-tailed macaque was used in retro-AAV tracing experiments. Animals were housed individually and kept on a 12-hour light/dark cycle. Food was provided daily with water *ad libitum*.

Environmental enrichment was provided daily, in the form of play objects or small food items. All procedures were approved by the Icahn School of Medicine IACUC and were carried out in accordance with NIH standards for work involving non-human primates.

### Virus prep

Modified sindbis virus for MAPseq was obtained from the MAPseq Core Facility at Cold Spring Harbor Laboratory^7^. The viral library used in this study had a diversity of 20,000,000 unique barcodes. Retro-AAV2 coding for mCherry (pAAV2-hSyn-mCherry, Addgene #114472, 2 x 10^13^ GC/ml) and green fluorescent protein (pAAV2-hSyn-eGFP, Addgene #50465, 2.2 x 10^13^ GC/ml) under the human synapsin promoter were obtained from Addgene. All viruses were stored at -80°C and aliquots were thawed over wet ice immediately prior to injection.

### Surgery and perfusion

#### Sindbis virus injections

For each animal, T1-weighted MRIs were obtained on a Skyra 3T scanner (Siemens) for surgical targeting. Animals were anesthetized with 5 mg/kg ketamine and 0.015 mg/kg dexmedetomidine; anesthesia was maintained with isoflurane as needed for the duration of the scan. Animals were scanned using an MRI-compatible stereotaxic frame (Jerry-Rig, Inc.). 3-4 images were obtained per scan, which were subsequently averaged together using the AFNI software suite (NIH)^53^. Then, stereotactic coordinates for the BLA could be computed using ImageJ (NIH)^54^. Two injection tracks were planned to target the middle of the amygdala’s anterior/posterior extent, equally spaced in the medial/lateral plane. One anterior injection track was also planned 1.5 mm in front of the middle of the amygdala, centered in the medial/lateral plane. Within each track, 4-5 injections 1.5 mm apart were planned in the dorsal/ventral plane to cover the entire extent of the basal and accessory amygdala nuclei.

After allowing approximately one week to elapse after the MRI, anesthesia was induced using ketamine and dexmedetomidine and maintained with isoflurane as described above. Surgery was performed under aseptic conditions, using the toothmarker method^55^ to place the animals in the same 3D position as the MRI. The skin, fascia, and muscles were retracted, and holes were drilled in the skull at each injection location using a surgical drill, widened if necessary, using rongeurs. Small dural incisions permitted Hamilton syringes access to the brain surface. 0.4 μl of virus was delivered at each injection site at a rate of 0.2 μl per minute, after which the needle was left in place for at least 3 minutes before reaching for the next site. Injections proceeded from the deepest site to the most superficial. The post-injection wait period was extended to at least 5 minutes at the top of each injection track before removing needles from the brain and proceeding to the next track.

After all injections were completed, the muscles, fascia, and skin were closed in anatomical layers. Following surgery, the animal was closely monitored in his home cage until normal behavior resumed. Postoperative treatment included buprenorphine (0.01 mg/kg, i.m., every 8 h) and meloxicam (0.2 mg/kg, i.m., every 24 h), based on attending veterinary guidance, as well as cefazolin (25 mg/kg, i.m., every 24 h) and dexamethasone sodium phosphate (0.4 –1 mg/kg, every 12–24 h) on a descending dose schedule.

Perfusion took place 67-72 hours after the final injection series. After being terminally anesthetized with ketamine/dexmedetomidine, the animal was prepared for RNAse-free perfusion. All tools were cleaned with RNAseZap solution (Fisher), and all solutions were prepared using RNAse-free reagents. The animal was perfused transcardially with ice-cold 1% paraformaldehyde (PFA; Electron Microscopy Science) in phosphate-buffered saline (PBS; Invitrogen) for approximately two minutes, followed by 4% PFA in PBS for approximately 18 minutes. Breathing was supplemented by manual ventilation until access to the heart was obtained.

Following brain extraction, the brain was placed in 4% PFA briefly before dissection and blocking as follows. After the cerebellum was removed, the brain was separated into hemispheres, the temporal lobes were dissected, and the remaining brain was cut into two blocks using a cryostat blade: one coronal cut was performed posterior to the central sulcus, separating the frontal and anterior parietal lobes from the remaining caudal portions of the brain – ensuring that the thalamus remained in the anterior block. Brain blocks were then post- fixed in 4% PFA for 48 hours. After post-fix, blocks were frozen slowly over dry ice before being stored at -80°C until sectioning.

#### Retro-AAV2 injections

Using the same MRI-guided stereotactic approach as above we targeted retro-AAV injections to the NAcc and AI in the left hemisphere and lOFC and vlPFC in the right hemisphere of a single monkey. Again, prior to surgery the animal was anesthetized, scanned at 3 T to obtain structural images, and injection targets were planned based on these scans. On the day of surgery, the animal was deeply anesthetized and injections were made during an aseptic neurosurgery. The skin, fascia, and muscles were opened in anatomical layers, burr holes were drilled over target locations and the dura opened. Injection syringes were then lowered into the brain and injections of retro-AAV2 were then made into the targets. At each location we injected 10 μl of virus, at a rate of 0.5 μl per minute. At the conclusion of each set of injections the needle was left in place for 5 minutes to allow the virus to diffuse. After all injections were completed, the muscles, fascia, and skin were closed in anatomical layers. Following surgery, the animal was closely monitored in their home cage until normal behavior resumed. Postoperative treatment included buprenorphine (0.01 mg/kg, i.m., every 8 h) and meloxicam (0.2 mg/kg, i.m., every 24 h), based on attending veterinary guidance, as well as cefazolin (25 mg/kg, i.m., every 24 h) and dexamethasone sodium phosphate (0.4 –1 mg/kg, every 12–24 h) on a descending dose schedule.

Eight weeks after surgery, the animals were perfused transcardially with ice-cold 1% PFA (Electron Microscopy Sciences) in PBS (Invitrogen) for approximately two minutes, followed by 4% PFA in PBS for approximately 18 minutes. The brains were then extracted, postfixed for 24 h in 4% PFA at 4°C and cryoprotected in 10% glycerol in PBS for 24 h, followed by 20% glycerol in PBS for another 24 h. The brains were then blocked with one coronal cut posterior to the thalamus, and the two resulting blocks were frozen in isopentane before storage at -80°C. The brain was then sectioned in the coronal plane on a freezing stage sliding microtome (Leica) at 40 µm in a 1:8 series. Tissue sets were stored either in PBS with 0.1% sodium azide (Sigma) at 4°C or in cryoprotectant comprised of glycerol, ethylene glycol, PBS, and distilled water (30/30/10/30 v/v/v/v, respectively) at -80°C.

#### Sectioning and dissection

From brains injected with sindbis virus, tissue was sectioned at 200 μm on a Leica 3050S cryostat that had been cleaned with RNAseZap prior to use. Sections were collected over dry ice and stored at -80°C prior to dissection. Cortical areas were then dissected according to sulcal landmarks over dry ice. The areas that were collected and our operational definitions of their boundaries can be found in this table.

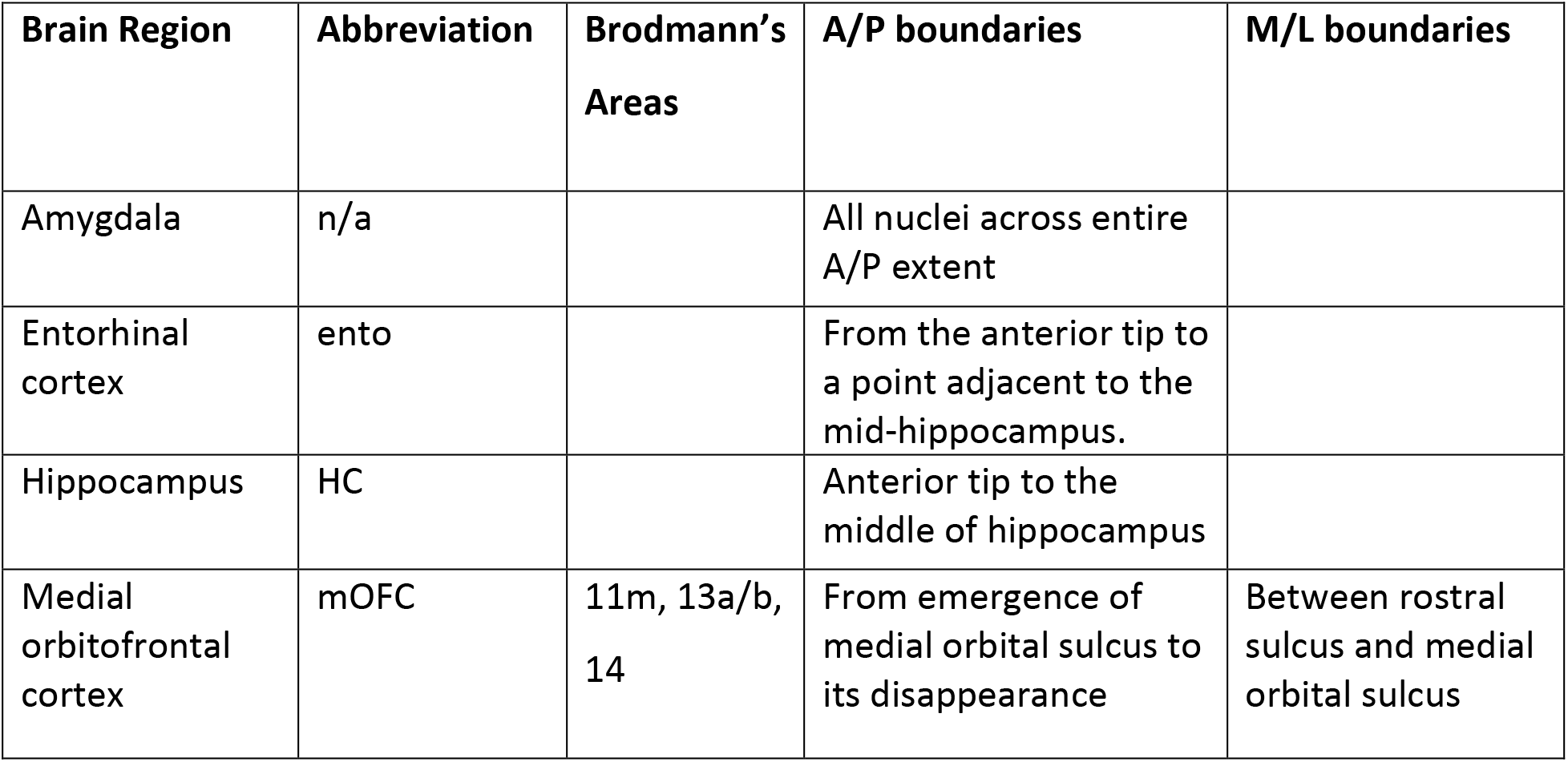
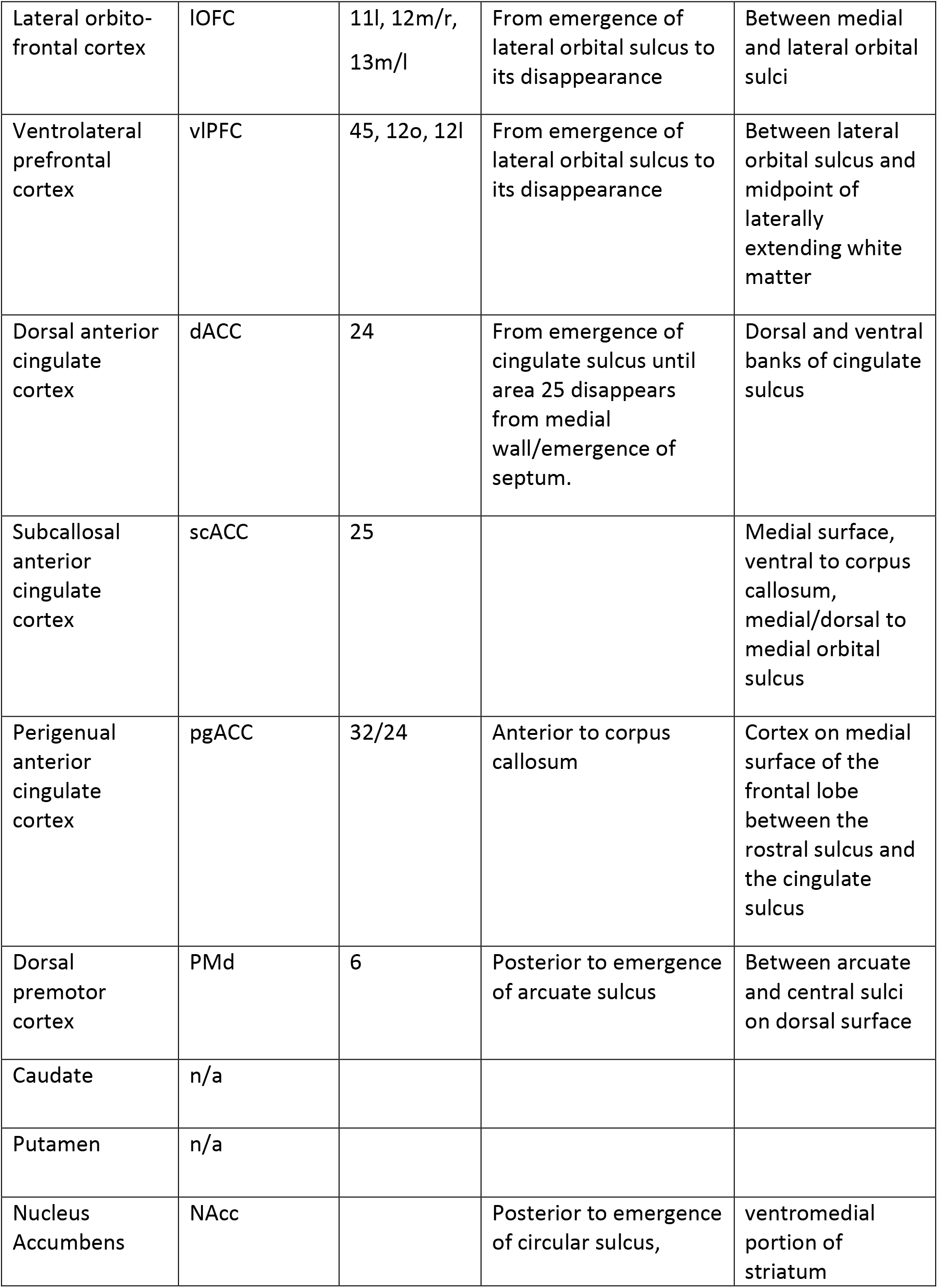
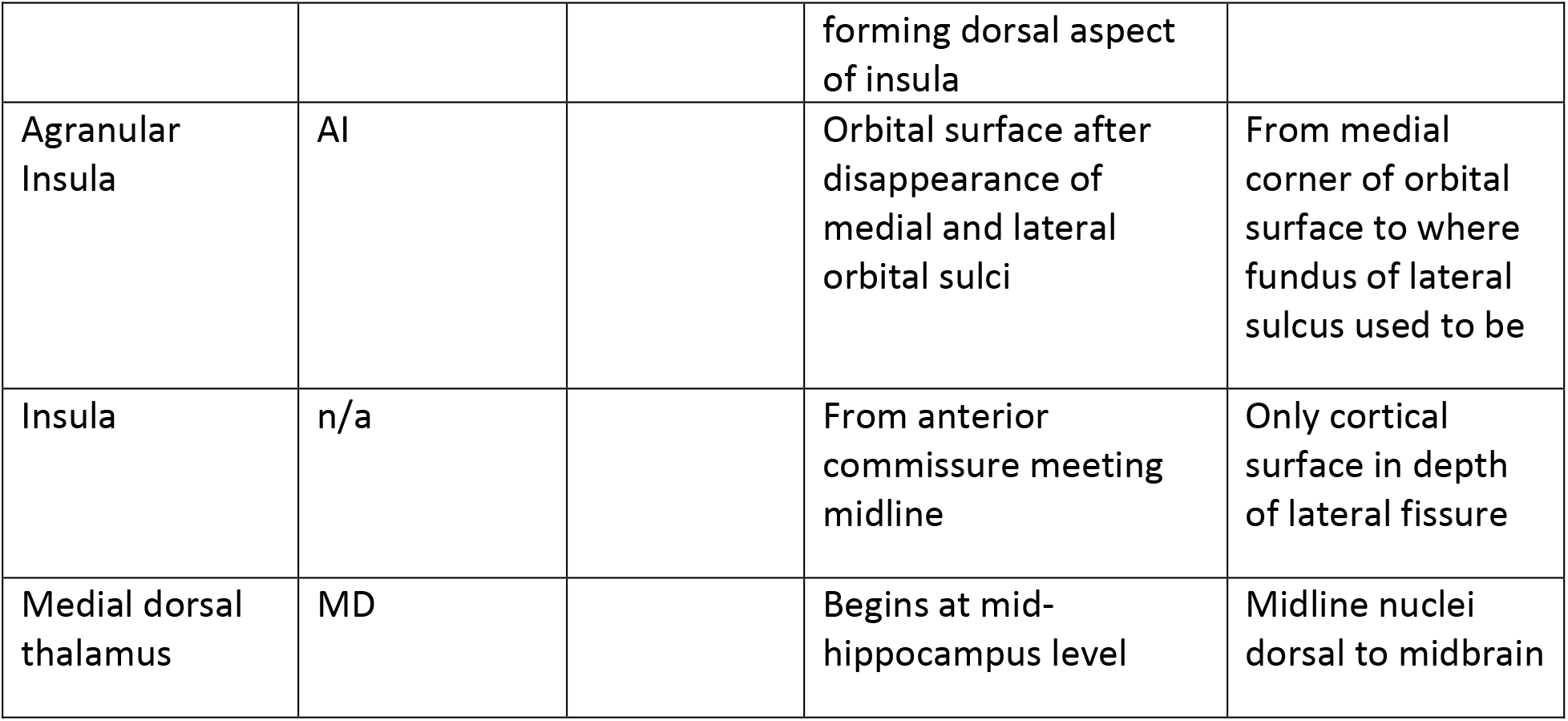

Samples from each area were combined across 3 sections in the anterior/posterior plane into 1.5-ml Eppendorf tubes, which were stored at -80°C prior to shipping frozen on dry ice for sequencing.

#### mRNA sequencing and preprocessing

Sequencing of MAPseq projections was performed by the MAPseq Core Facility at CSHL as described in Kebschull et al. 2016^7^. Briefly, sections were homogenized and treated with protease before RNA extraction. Total RNA was extracted using an established Trizol-based protocol. RNA quality was verified on Bioanalyzer and bulk amounts of barcodes were examined by qPCR and compared with the amount of housekeeping gene prior to sequencing. Barcode RNA was reverse transcribed into cDNA, and a known amount of RNA (spike-in sequence) was added to each sample during reverse transcription. Barcode cDNA was then double-stranded, PCR amplified to produce a sequencing library, and the purified barcode library was then submitted for a paired-end36 run on an Illumina NextSeq machine.

Preprocessing was performed by CSHL using MATLAB (Mathworks) to create a barcode matrix with size (n samples x n barcodes). Corresponding barcode sequences and spike-in counts are also extracted to allow for normalization and assessment of duplicates.

#### Filtering and analysis

Filtering was performed in Python 3.9^56–59^ using a modification of the publicly available normBCmat.m script^7^; specific analyses can be found in the code available on Github. Briefly, for each animal, barcodes are filtered, then the raw barcode counts are divided by the spike-in counts for normalization; barcodes survive filtering if the max barcode amount is found in the injection sites (amygdala), the max count is greater than 20, and if any other sample (whether within amygdala or outside) has greater than 5 barcode counts. This adaptation from the original filtering allows barcodes which send their strongest projections only within amygdala to survive filtering.

The resulting barcode matrix was then binarized and collapsed within brain regions: if a barcode was found in any of the multiple samples for each brain area, then we counted that neuron as projecting to that brain area. This collapsing and binarization allowed us to combine the results from the four sequencing runs: each of the two hemispheres from one animal, and one hemisphere sequenced twice from the other animal. Only unique barcodes from the re- sequencing were included in analyses; duplicates were removed based on their barcode sequences. This filtered, binarized, and collapsed matrix combined across the four hemispheres was the basis for all analyses in this study.

#### Population summary

Projection proportions were calculated by summing the barcodes found in one brain area and dividing by the total number of barcodes in the dataset; chance level was defined as the barcode proportion found in control sites collected from cerebellum. The number of targets for each neuron is calculated by summing the number of areas in which individual barcodes were found and subtracting one, to count the number of non-amygdala targets for each neuron; thus, a neuron which projects from one amygdala sample to another was defined as having zero targets.

#### Conditional probabilities

To calculate the conditional probability of a neuron projecting to two areas, A and B, we first found all of the cells that projected to area A (irrespective of their other targets). Then, we found the subset of those cells which also project to area B. Thus, the conditional probability of B|A is the number of cells that project to both A and B divided by the number of cells that project to A.

#### K-means clustering

We performed k-means clustering using *scikit-learn* Python function. The optimal number of clusters was determined using the elbow method, in which we plotted the number of clusters against the within-cluster sum of squares. We found k= 12 to be optimal. We then sorted the projection matrices by k-means cluster.

#### Over- and under-represented motifs

To assess whether any branching motifs were under- or over- represented compared to chance, we first constructed a null distribution based on the overall proportion of barcode found in each area. We assumed that the probability of a neuron branching to two areas, A and B, would be the product of the independent probabilities of projecting to A and B: *p*(*A&B*) = *p*(*A*) ∗ *p*(*B*). This approach was taken for bi-, tri-, and quadrifurcations. We did not pursue analysis of neurons with five or more targets, as there were fewer than 100 neurons with each number of targets, leading to insufficient sample size on which to perform count-based statistics.

Then, we compared our actual barcode counts to the expected null distribution using a binomial probability test (*scipy.stats.binomtest*). Resulting p-values were FDR-corrected within each branching degree, and effect sizes were calculated as log_!_(^"#$%&’%(^).

#### Network analysis

To compare projections within the medial and ventral frontal cortex, we first isolated projections to only one, not multiple, of the areas. We defined these networks by comparing areas on the orbital surface (mOFC, lOFC, vlPFC, AI) to one another and areas on the medial surface (dACC, pgACC, scACC). First, we computed the degree of overlap within these networks by preparing Venn diagrams of projections specific to the areas and which branched between them. The proportion of branching within each area was compared using z-tests for proportions; p-values were adjusted using FDR correction.

We then focused on the projections which were specific while excluding any neurons which projected to multiple areas in the same network. Projection strength to other areas were calculated as described above and compared using pairwise z-tests for proportions and Fisher’s exact tests, corrected for multiple comparisons. Degrees of branching were calculated as before, and branching was compared between brain areas using a permutation test in which area labels were shuffled 1000x to generate a null distribution; distributions were then compared using Chi-squared tests. Clustered projection heatmaps were constructed using *seaborn.clustermap* with Ward’s distance.

#### Comparison of neurons across populations

To understand whether projection motifs were reproducible across hemispheres and across animals, we compared each of our independent sequencing runs to simulated neurons from a uniform distribution (Extended Data Figure 4). 250 neurons were randomly sampled from each of the sequencing runs and compared to other runs and to the simulated neurons using cosine distance.

#### Stereology

Tissue for stereology was mounted from PBS onto gelatin coated slides and mounted and coverslipped using Vectashield Vibrance Antifade aqueous mounting medium (Vector Labs). Slides were stored in a lightproof slide box at 4°C to prevent fluorophore fading during analysis.

An adjacent series was histochemically stained for acetylcholinesterase (AChE) for more reliable identification of amygdala boundaries. Tissue was incubated overnight at 4°C in a solution containing 0.68% sodium acetate (Thermo Scientific), 0.1% copper (II) sulfate (Thermo Scientific), 0.12% glycine (Thermo Scientific), 0.12% acetylthiocholine iodide (TCI America), and 0.003% ethopropazine (Sigma Aldrich). The following morning, sections were rinsed 3x in PBS for 5 minutes each, transferred to a solution of 0.1 M acetic acid (LabChem) with 1% sodium sulfide (Thermo Scientific) for 1-2 minutes, and then rinsed again 3x with PBS before mounting as above.

Stereology was performed using a Zeiss Apotome.2 microscope equipped with a Q- Imaging digital camera, motorized stage, and Stereo Investigator software (MBF Bioscience). A total of 12 sections were used for stereology, evenly distributed to cover the anterior-posterior extent of the amygdala. The borders of the amygdala were identified using a 5x objective on the AChE-stained sections, then ROI contours were realigned with the unstained sections for stereological analysis. EGFP-, mCherry-, and double-labelled cells were counted based on the soma as the counting target; the optical fractionator probe was used for stereological estimation as described in West et al., 1991^38^. Neurons were counted under a 10x objective, with a counting frame of size 150 x 150 x 15 μm; a 5-μm guard was applied to the dorsal aspect of each section and a 20-μm guard to the ventral side. Counting frames were arranged in a 670.8 x 670.8 μm grid for systematic-random sampling. Results of the stereological analysis were compared using z-tests for proportions.

#### fMRI analysis

Functional connectivity analyses were conducted on a previously published resting-state functional MRI dataset^41^. We re-analyzed a subset of the scans, including data from 6 rhesus macaques (5 males), focusing only on the control and pre-injection scans. After performing standard preprocessing and warping all the brains to a standard template, masks for target areas were drawn according to the areal boundaries used for MAPseq utilizing the D99, CHARM, and SARM atlases as a basis^60–63^. Then, functional connectivity was computed between each amygdala voxel and each target voxel in the ipsilateral hemisphere^64^. Correlations were z-transformed, and all animals’ data was combined into one large dataset after computing within-animal z-scores. This way, we could analyze all of the amygdala voxels from each animal without averaging across animals. The following analyses utilized the voxel-wise connectivity matrix. Whole brain functional connectivity for the amygdala was calculated by averaging the time-series within the anatomically defined amygdala and then correlating the average time series with every voxel within the brain, for each animal, before averaging across animals.

The continuous measure of functional connectivity was binarized as follows. For each amygdala voxel, the peak target connectivity was found; other target voxels which had connectivity z-scores within 70% of that maximum were said to be connected, while those under that threshold were said to not be connected. That 70% threshold allows about 5% of the target voxels to survive filtering (**Supplemental Figure 6**). Then, as in MAPseq, we collapsed the data across voxels within each target area, such that if one voxel in a target area survived filtering, we concluded that amygdala was ‘functionally connected’ to that area. Then, we followed a similar analysis pipeline as for the MAPseq data; computing projection strengths by counting the proportion of voxels in each area to have survived filtering, computing conditional probabilities, counting the number of ‘targets’ for each voxel, and performing k-means clustering. We also performed the same network analysis by filtering for voxels connected to areas in ventral or medial frontal cortex.

## Data and Code Availability

Data and code can be made available upon reasonable request by reaching out to.

## Extended Data Figures

**Extended Data Figure 1:**
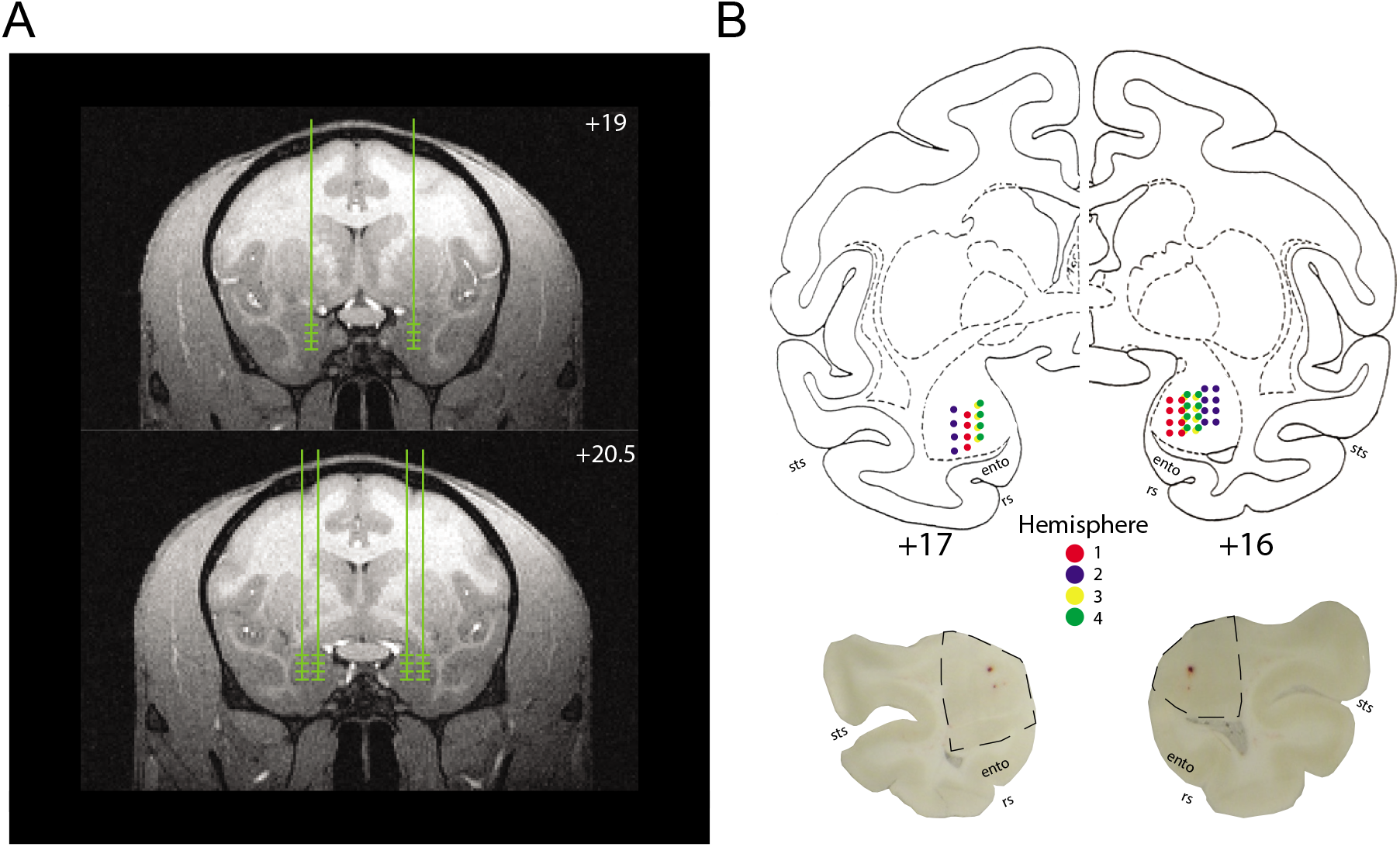
Anatomical verification. **A**) Representative MRI images showing anterior (top) and middle (bottom) injection targets within amygdala. Vertical lines indicate intended injection tracks, while horizontal lines indicate injection depths along those tracks. **B**) Locations of injections for individual animals (colors); anterior injection on the left, middle on the right. Example tissue sections shown below with the extent of the amygdala surrounded by the dotted line (ento refers to entorhinal cortex, rs rhinal sulcus, sts superior temporal sulcus).

**Extended Data Figure 2.**
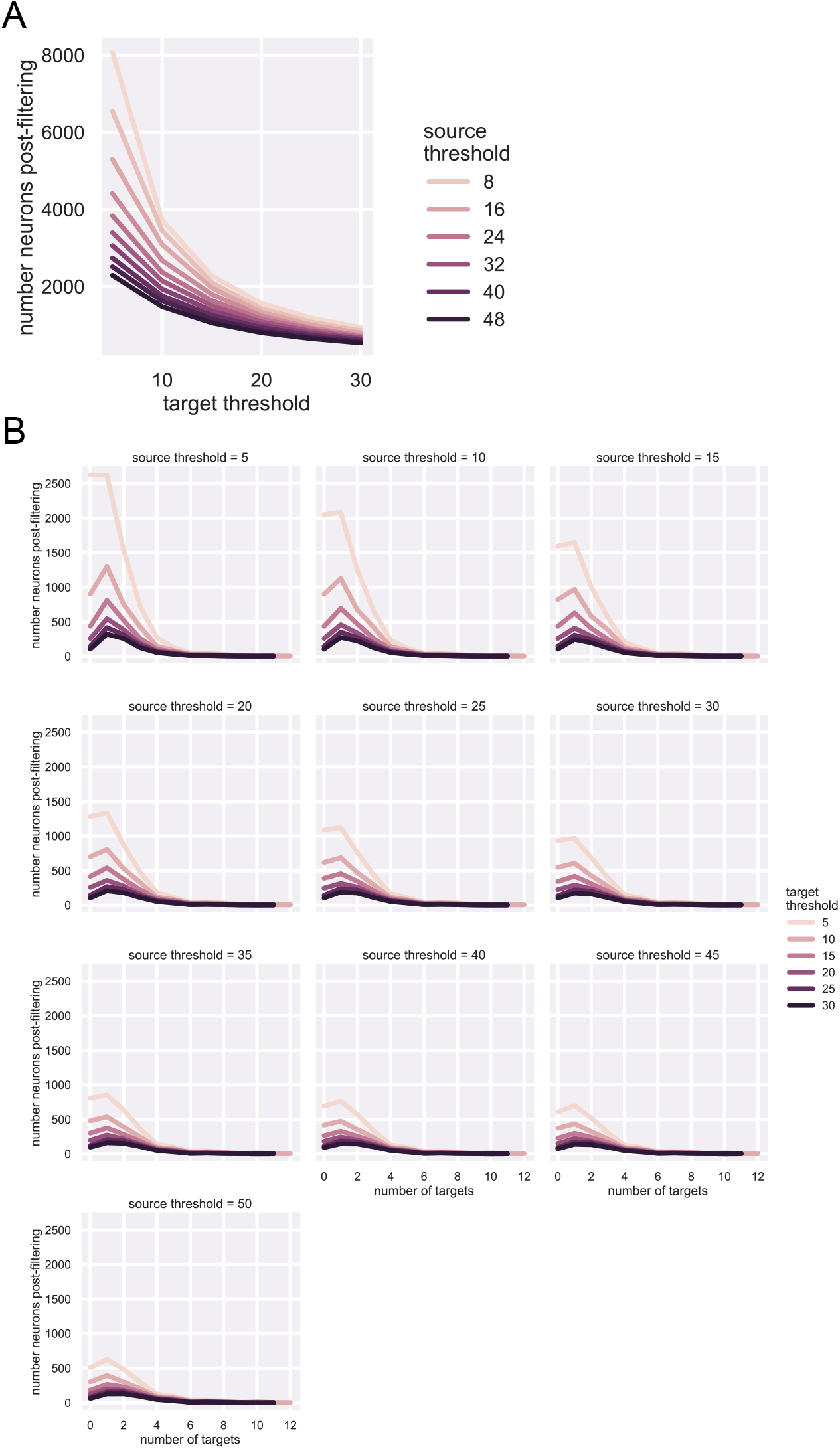
Filtering parameters do not dramatically change recovered barcodes. **A**) Number of barcodes surviving filtering for different source thresholds (color of line) and target thresholds (x-axis). Sufficient source threshold eliminates majority of noise. **B**) Effect of thresholding on number of projection targets per neuron. Each plot is one source threshold, while colored lines reflect different target thresholds. The shape of the distributions is lightly flattened by increasing projection threshold, while again, source threshold is responsible for most of the noise.

**Extended Data Figure 3:**
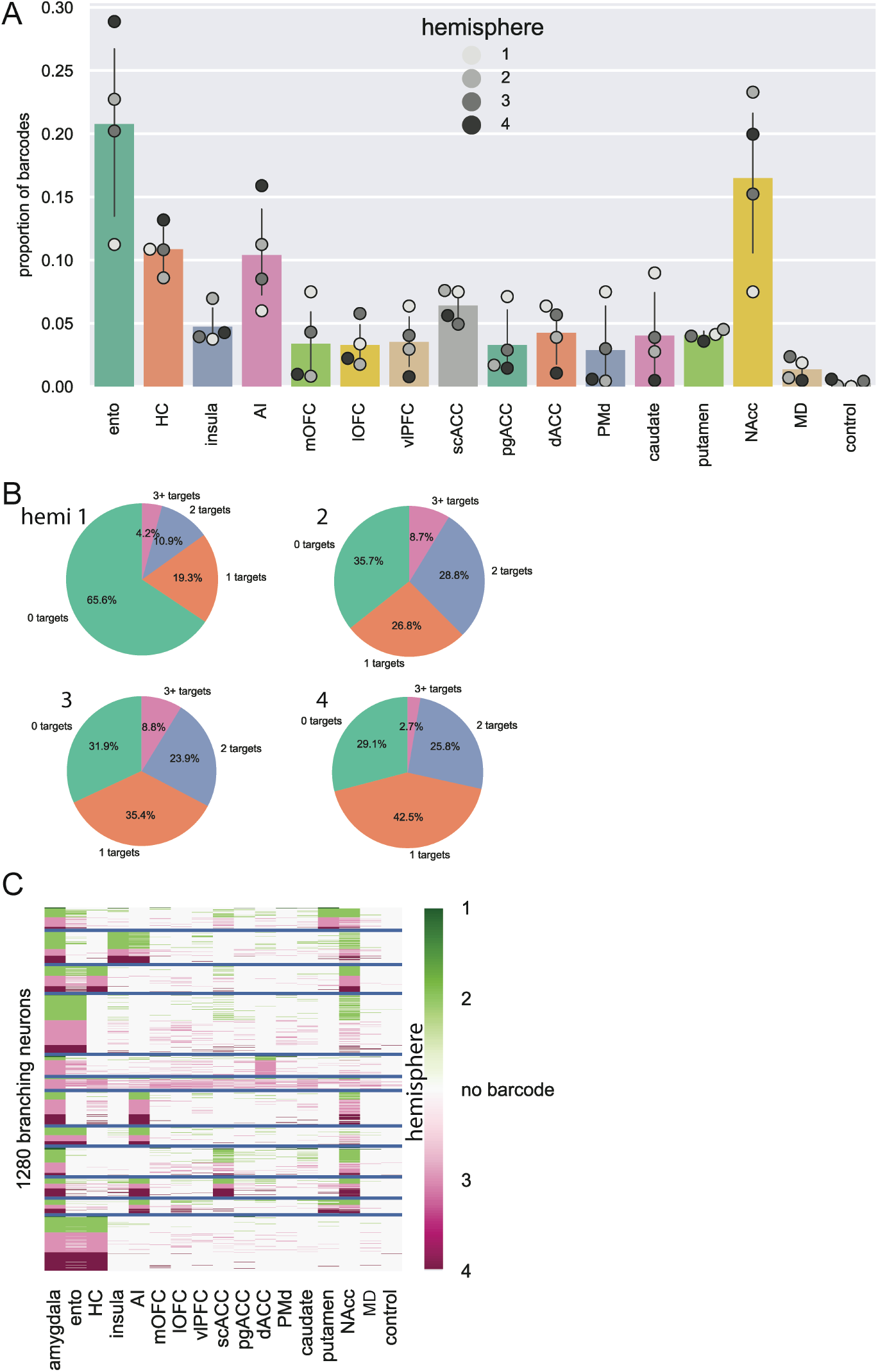
MAPseq is consistent across animals. **A**) Overall barcode distribution across areas. Colored bars represent the mean across all 4 hemispheres sequenced, error bars are standard deviation, and individual points reflect counts within each hemisphere’s data separately. **B**) Number of targets for each neuron across hemispheres – roughly equal proportions of 1-target and 2+-target neurons, with most variance observed in proportion of 0- target neurons. **C**) K-means clustered branching projections, labelled by hemisphere. Note that most clusters are comprised of neurons from multiple hemispheres.

**Extended Data Figure 4:**
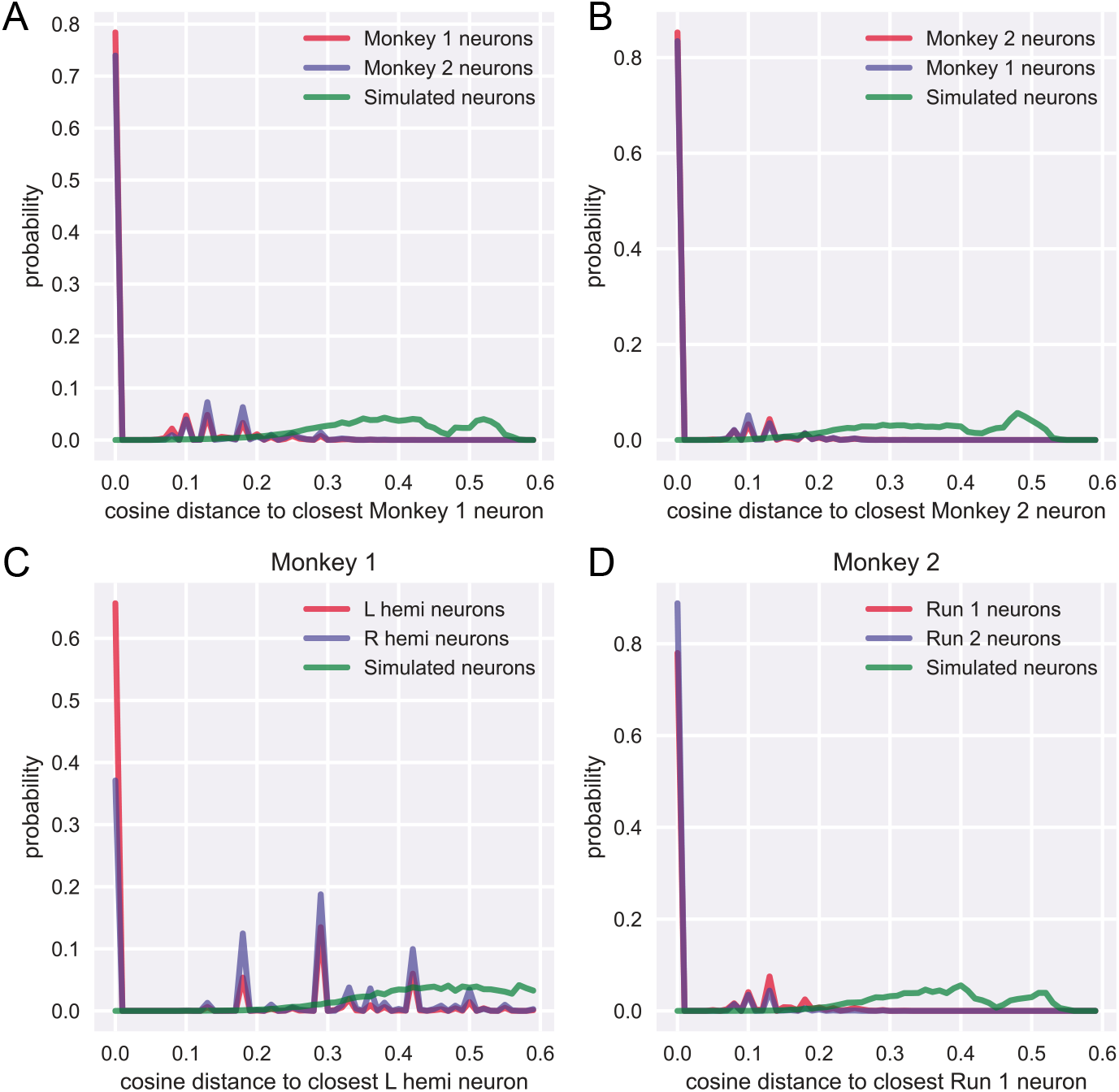
Comparison across hemispheres. Density plot of cosine distance between two actual samples (red/blue) and simulated neurons from a uniform distribution (green). **A**) Monkey 1 neurons (both hemispheres) as a basis are more similar to monkey 2’s neurons (Kolmogorov-Smirnov test, D = 0.12, *p* = 0.81) than the simulated neurons (D = 0.52, p < 0.0001. **B**) Same for monkey 2 as a basis. **C**) Within monkey 1, the two hemispheres are more similar to each other (D = 0.18, *p* = 0.27) than the random neurons (D = 0.43, *p* < 0.0001). **D**) Within monkey 2, the two sequencing runs were more similar to each other (D = 0.22, *p* = 0.12) than the simulated neurons (D = 0.48, *p* < 0.0001).

**Extended Data Figure 5:**
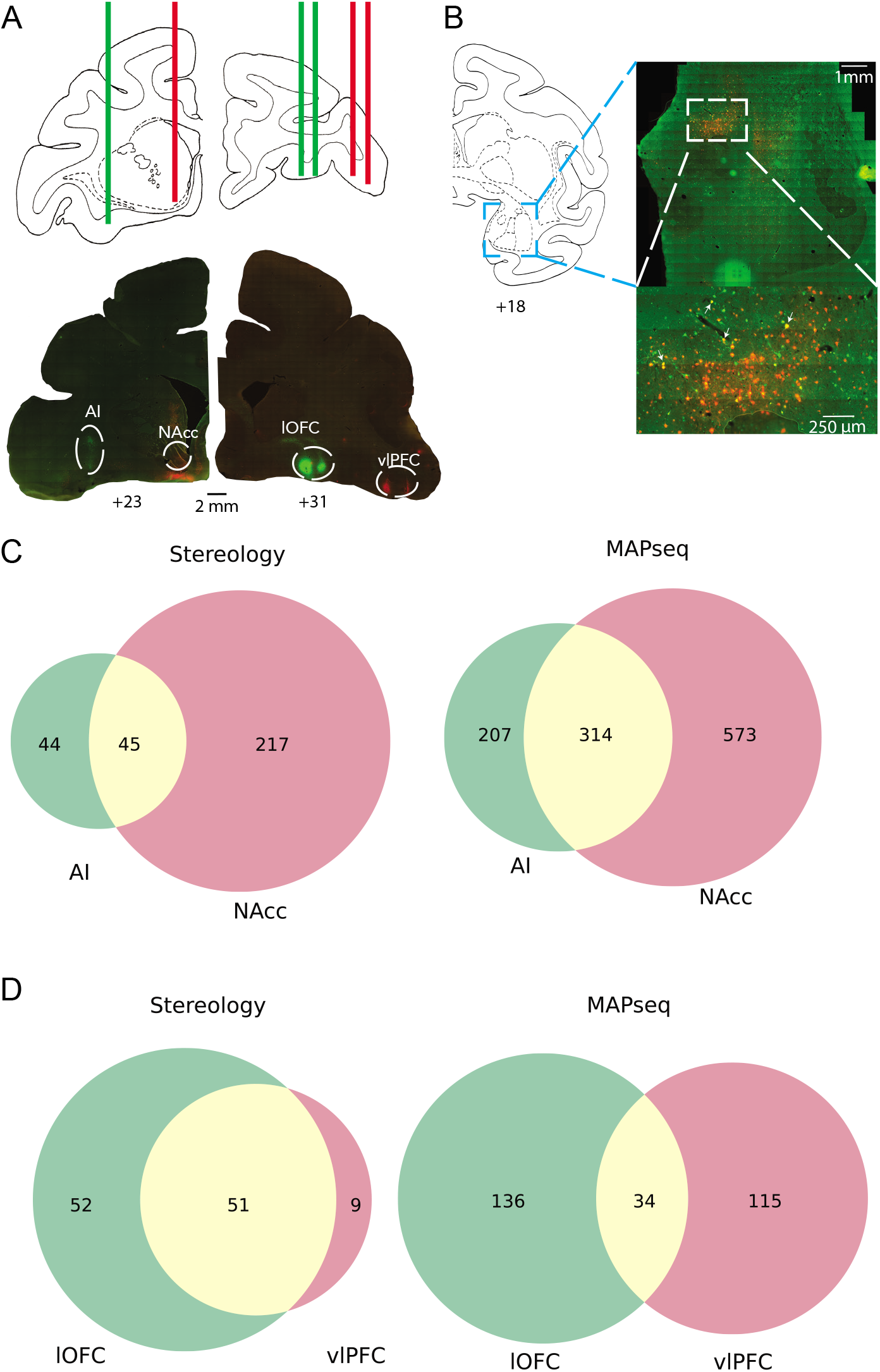
Stereological confirmation of branching motifs. **A**) Top: injection strategy plotted on atlas sections (distances from intra-aural plane in mm); red and green lines refer to mCherry and EGFP retro-AAVs. Bottom: photomicrographs of actual injections sites. **B**) Example retrograde labelling in amygdala (shown in atlas on right). The bottom image is comprised of approximately 4x4 tiled 10x magnified images. Double labelled cells are non-exhaustively labelled with arrows. **C**) Stereology results (left) from left hemisphere injections compared to MAPseq results (right). **D**) Same for right hemisphere injections.

**Extended Data Figure 6:**
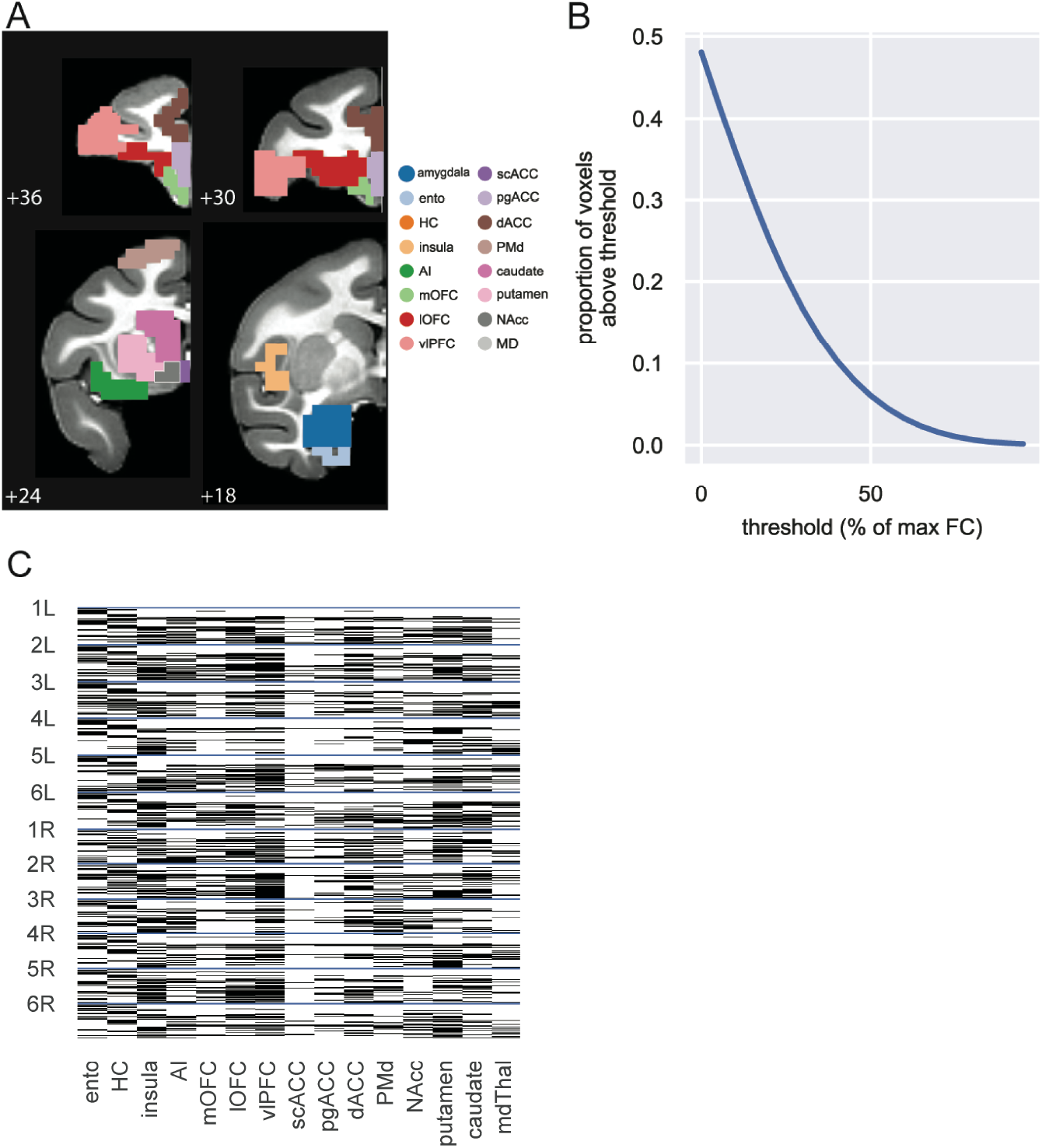
fMRI analysis. **A**) ROIs used for fMRI analysis shown on NMT atlas slices (hippocampus and MD not shown). **B**) Proportion of voxels determined to be ‘functionally connected’ decreases with increasing threshold (the threshold was set at 70% because ∼5% of voxels survive filtering). B) Binarized and collapsed MRI connectivity. Number labels refer to individual animals, while L and R refer to left and right hemispheres, respectively; only ipsilateral connections were assessed.

