## Supplemental figures only for "High-throughput sequencing of macaque basolateral amygdala projections reveals dissociable connectional motifs with frontal cortex"

### Extended Data Figures

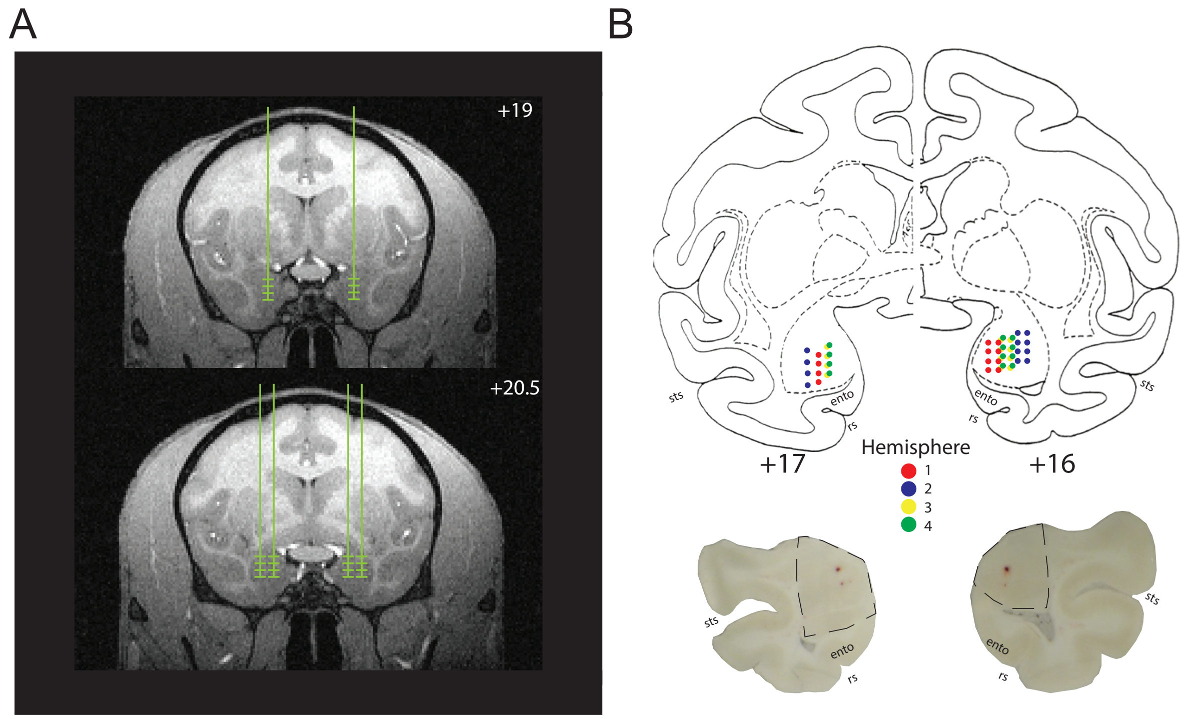


**Extended Data Figure 1: Anatomical verification. A**) Representative MRI images showing anterior (top) and middle (bottom) injection targets within amygdala. Vertical lines indicate intended injection tracks, while horizontal lines indicate injection depths along those tracks.
**B**) Locations of injections for individual animals (colors); anterior injection on the left, middle on the right. Example tissue sections shown below with the extent of the amygdala surrounded by the dotted line (ento refers to entorhinal cortex, rs rhinal sulcus, sts superior temporal sulcus).

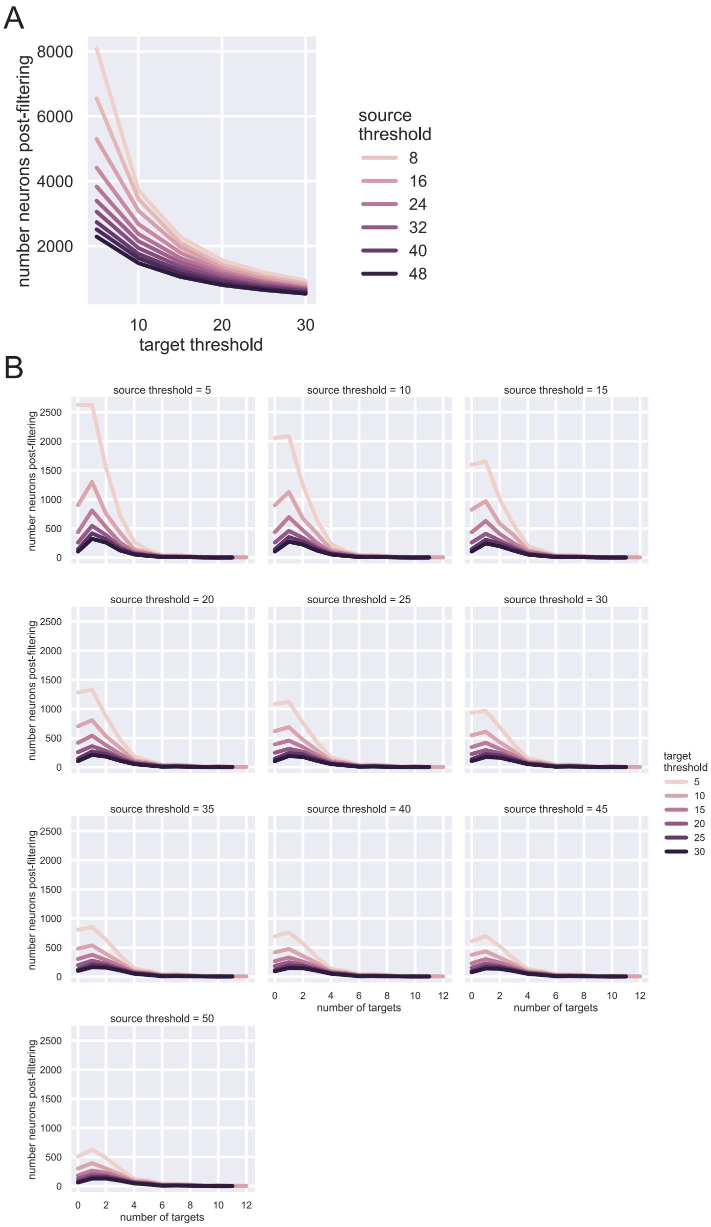


**Extended Data Figure 2. Filtering parameters do not dramatically change recovered barcodes.** **A**) Number of barcodes surviving filtering for different source thresholds (color of line) and target thresholds (x-axis). Sufficient source threshold eliminates majority of noise. **B**) Effect of thresholding on number of projection targets per neuron. Each plot is one source threshold, while colored lines reflect different target thresholds. The shape of the distributions is lightly flattened by increasing projection threshold, while again, source threshold is responsible for most of the noise.

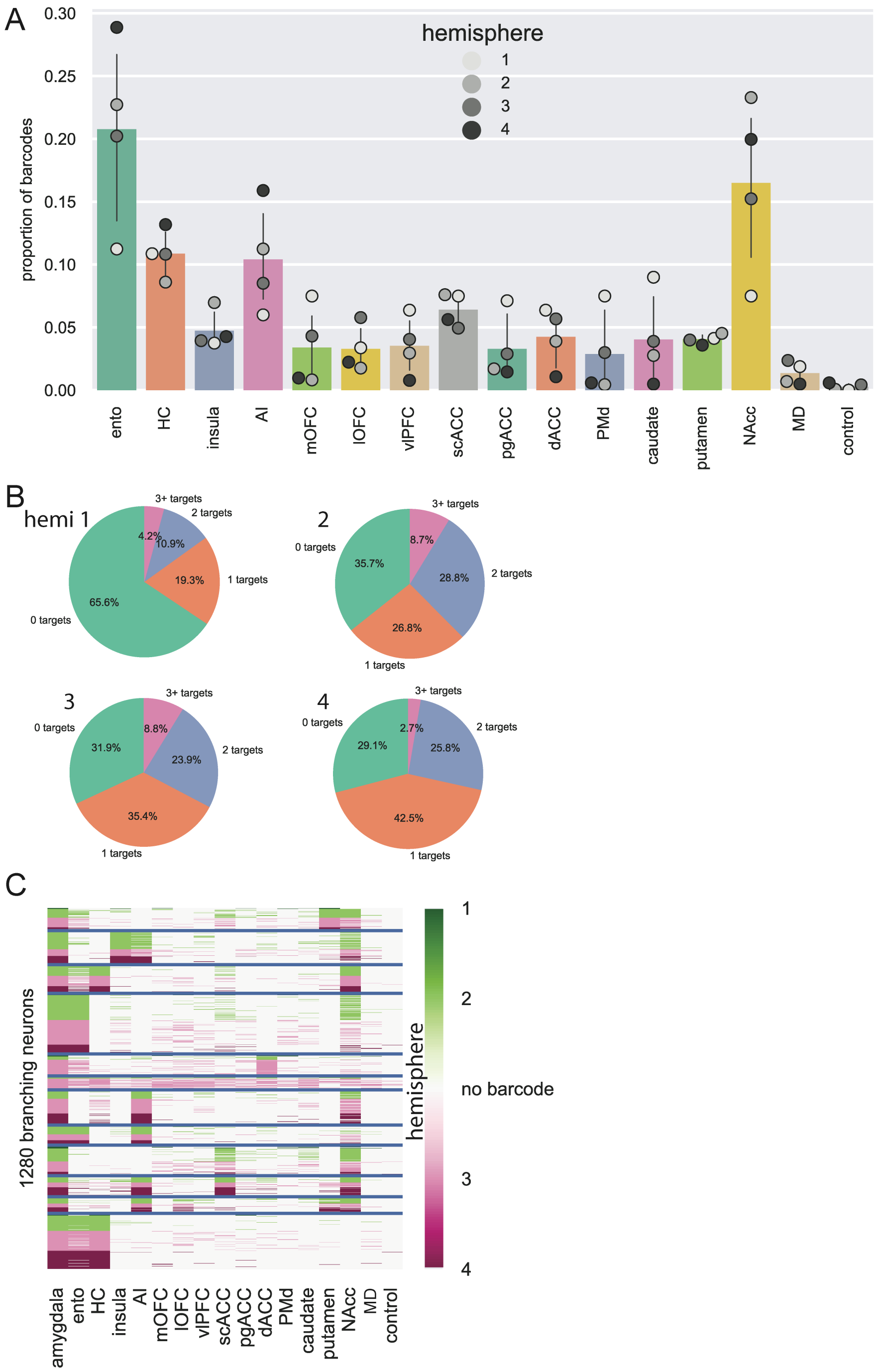


**Extended Data Figure 3: MAPseq is consistent across animals. A**) Overall barcode distribution across areas. Colored bars represent the mean across all 4 hemispheres sequenced, error bars are standard deviation, and individual points reflect counts within each hemisphere's data separately. **B**) Number of targets for each neuron across hemispheres – roughly equal proportions of 1-target and 2+-target neurons, with most variance observed in proportion of 0-target neurons. **C**) K-means clustered branching projections, labelled by hemisphere. Note that most clusters are comprised of neurons from multiple hemispheres.

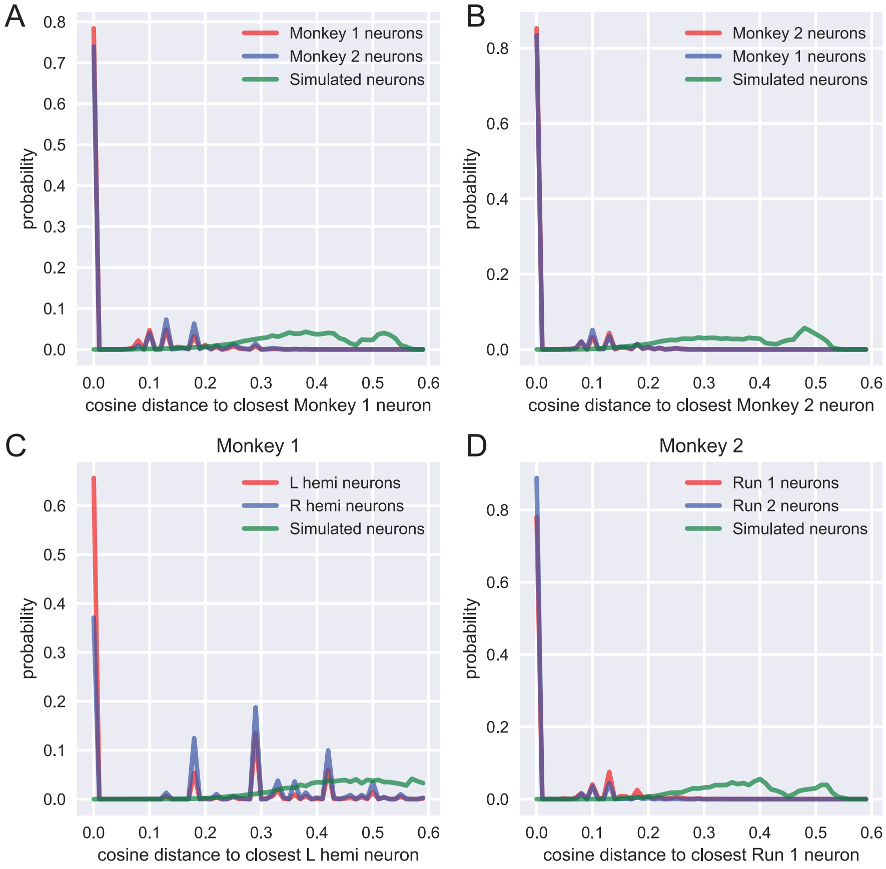


**Extended Data Figure 4: Comparison across hemispheres.** Density plot of cosine distance between two actual samples (red/blue) and simulated neurons from a uniform distribution (green). **A**) Monkey 1 neurons (both hemispheres) as a basis are more similar to monkey 2's neurons (Kolmogorov-Smirnov test, D = 0.12, $p = 0.81$) than the simulated neurons (D = 0.52, p < 0.0001. **B**) Same for monkey 2 as a basis. **C**) Within monkey 1, the two hemispheres are more similar to each other (D = 0.18, $p = 0.27$) than the random neurons (D = 0.43, $p < 0.0001$). **D**) Within monkey 2, the two sequencing runs were more similar to each other (D = 0.22, $p = 0.12$) than the simulated neurons (D = 0.48, $p < 0.0001$).

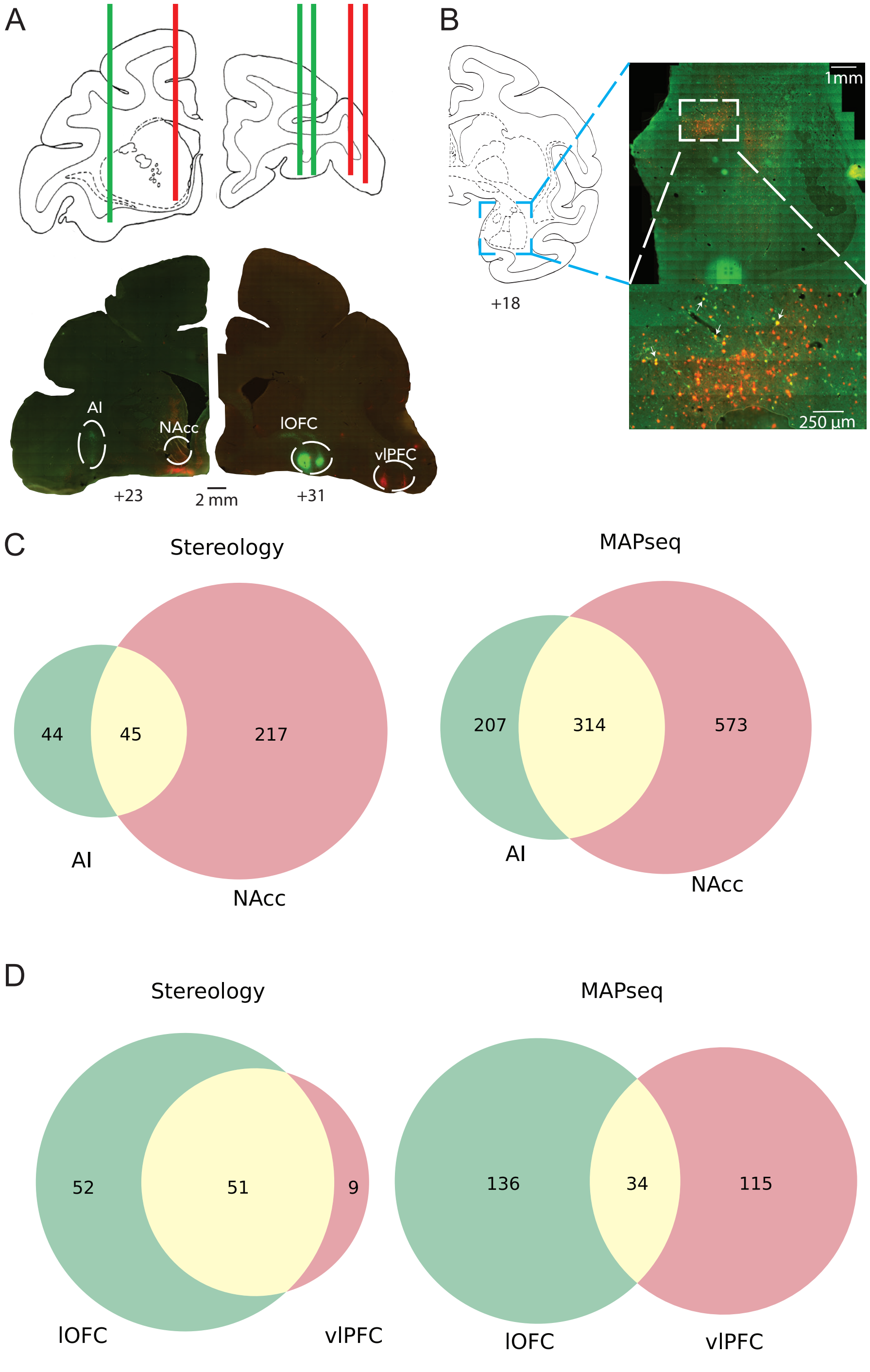


**Extended Data Figure 5: Stereological confirmation of branching motifs. A**) Top: injection strategy plotted on atlas sections (distances from intra-aural plane in mm); red and green lines refer to mCherry and EGFP retro-AAVs. Bottom: photomicrographs of actual injections sites. **B**) Example retrograde labelling in amygdala (shown in atlas on right). The bottom image is comprised of approximately 4x4 tiled 10x magnified images. Double labelled cells are non-exhaustively labelled with arrows. **C**) Stereology results (left) from left hemisphere injections compared to MAPseq results (right). **D**) Same for right hemisphere injections.

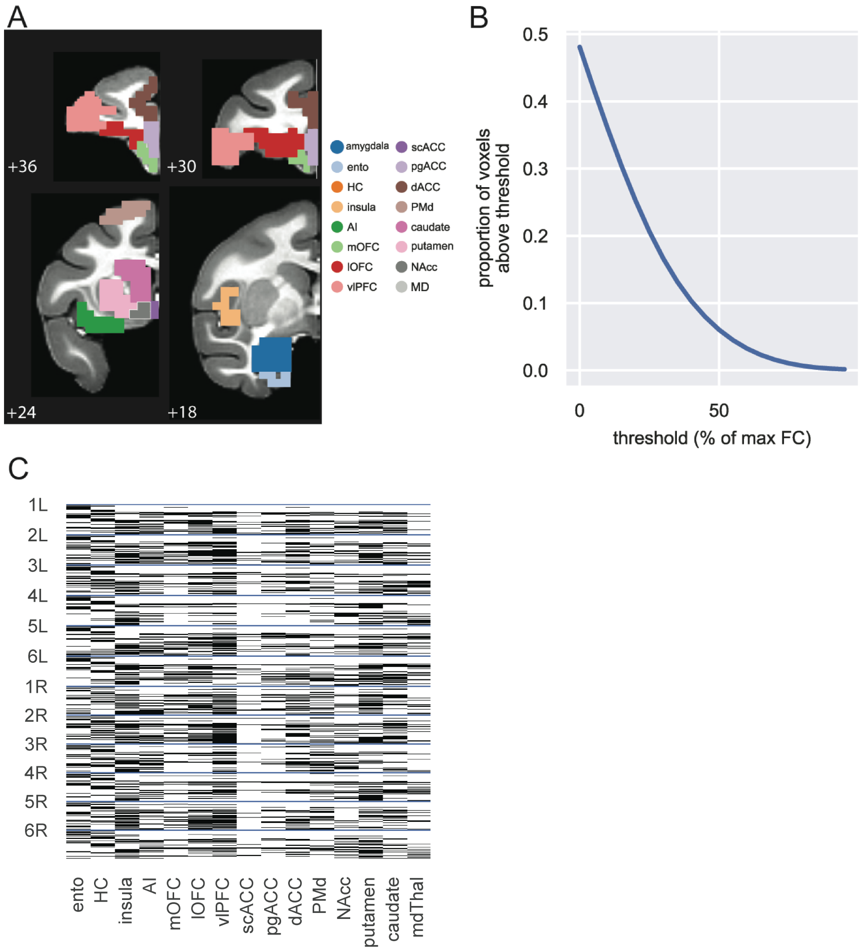


**Extended Data Figure 6: fMRI analysis. A**) ROIs used for fMRI analysis shown on NMT atlas slices (hippocampus and MD not shown). **B**) Proportion of voxels determined to be 'functionally connected' decreases with increasing threshold (the threshold was set at 70% because ~5% of voxels survive filtering). B) Binarized and collapsed MRI connectivity. Number labels refer to individual animals, while L and R refer to left and right hemispheres, respectively; only ipsilateral connections were assessed.
